# Metabolism of trichloroethylene following inhalation exposure in mice

**DOI:** 10.64898/2026.10.13.758615

**Authors:** Jocelyn Dicent, Carolina Duarte-Hospital, Neda M. Ilieva, Randolph R. Singh, Briana R. De Miranda, Gary W. Miller

## Abstract

Trichloroethylene (TCE) is a pervasive environmental pollutant associated with elevated risk of developing Parkinson’s disease (PD). Studies characterizing TCE metabolism in humans indicate that 40–75% of the absorbed dose is metabolized, suggesting that TCE toxicity may be attributable to its metabolites. However, there is limited knowledge of TCE metabolite neurotoxicity. We conducted a TCE inhalation study in mice followed by high-resolution mass spectrometry (HRMS) on brain, liver, and blood to identify which TCE metabolites were present following human-relevant TCE exposure. Mice were exposed to 100 ppm TCE for 4 hours. Plasma, brain, and liver were collected from each animal up to 24 hours after exposure. We developed a method capable of detecting TCE metabolites trichloroacetic acid (TCA), dichloroacetic acid (DCA), monochloroacetic acid (MCA), trichloroethanol (TCEOH), trichloroethanol glucuronide (TCEOH-gluc), S-(1,2-dichlorovinyl)glutathione (DCVG), S-(1,2-dichlorovinyl)-L-cysteine (DCVC), *N*-acetyl-*S*-(1,2-dichlorovinyl)-L-cysteine (NAcDCVC), and untargeted metabolites using a single extraction protocol on a liquid chromatography-based mass spectrometry system (LC-MS/MS). Targeted analysis of TCE-exposed biological samples identified DCA, TCA, and TCEOH-gluc. Untargeted analysis of TCE-exposed mouse brain revealed previously established and novel pathway perturbations associated with TCE exposure. Together, these findings indicate that DCA, TCA, and TCEOH-gluc are the most prevalent TCE metabolites in mice. The novel pathways perturbed by TCE exposure warrant further exploration to better understand the mechanisms underlying the toxicity of TCE and its metabolites.

## Introduction

Trichloroethylene (TCE) is an environmentally pervasive halocarbon solvent, with known exposure primarily occurring through drinking water or vapor intrusion (Wu and Schaum 2000; U.S. Environmental Protection Agency [EPA] 2025). Growing evidence of TCE toxicity has led to restrictions on its use as an industrial degreaser and dry-cleaning compound in the U.S. (Smith 1966; Agency for Toxic Substances and Disease Registry [ATSDR] 2019; U.S. Environmental Protection Agency [EPA] 2023). TCE exposure has been increasingly linked to Parkinson’s Disease (PD), with experimental evidence aligning with existing literature on the predominant molecular mechanisms perturbed in PD (Guehl et al. 2003; Gash et al. 2008; Liu et al. 2010a; Goldman et al. 2012; Agency for Toxic Substances and Disease Registry 2017; P.C. Keane et al. 2019; De Miranda et al. 2021; Ilieva et al. 2022; Goldman et al. 2023; Adamson et al. 2023). While the TCE-PD association is strong, it remains to be determined whether the neurotoxic effects of TCE are driven by the parent compound or its metabolites (Greenamyre and Hastings 2004; Subramaniam and Chesselet 2013; MacMahon Copas et al. 2021). We have yet to understand whether those metabolites are present in the brain, either through onsite formation of metabolites or circulation, following a human-relevant exposure. Understanding whether these metabolites are present in the brain is critical to understanding their neurotoxicity.

Upon exposure to TCE, 40 – 75% of the compound is metabolized, indicating that examining TCE metabolites may be critical to understanding its toxic effects (Lash et al. 2000; Bonvallot et al. 2010). TCE undergoes biotransformation via two principal routes: the cytochrome P450 2E1 (CYP2E1) pathway, which generates rapidly excreted, water-soluble metabolites, and the glutathione S-transferase (GST) pathway, which generates metabolites that are typically associated with the toxification of TCE (Lash et al. 2000; Bonvallot et al. 2010; De Miranda and Timothy Greenamyre 2020). Some TCE metabolites, such as trichloroacetic acid (TCA), have longer half-lives than TCE itself, suggesting a longer-lasting effect than the parent compound (Zimmermann et al. 1998; International Agency for Research on Cancer [IARC] 2004). Additionally, many CYP2E1 metabolites appear in urine within minutes of inhalation or oral exposure to TCE, further suggesting that these metabolites may contribute to disease outcomes (Souček and Vlachová 1960; Lash et al. 2000).

Given its lipophilic nature, TCE readily crosses membranes including the blood-brain barrier. Information on the propensity of its metabolites to cross the blood-brain barrier is limited to 1-trichloromethyl-1,2,3,4-tetrahydro-β-carboline (TaClo) (Riederer et al. 2002; Cichocki et al. 2016; P. C. Keane et al. 2019; Liu et al. 2025). Through inhalation, TCE may bypass the blood-brain barrier entirely by migrating to the brain via the olfactory nerve, suggesting that metabolites detected in the brain may reflect local rather than hepatic metabolism (Lucchini et al. 2012; Jedličková et al. 2025). Currently, it remains unclear whether TCE neurotoxicity via inhalation is mediated by its metabolites, given limited information on their presence in the brain following exposure. Identifying which metabolites are present in the brain is a necessary step toward understanding their role in neurodegeneration.

Current TCE metabolite detection methods are resource-intensive and laborious, typically requiring multiple sample preparations, depending on the compound class. (Tanaka and Ikeda 1968; Lash et al. 1999; Song and Ho 2003; Delinsky et al. 2005; Luo, Furuya, et al. 2018; Luo, Hsieh, et al. 2018a) These methods are promising, but they may be difficult to apply in studies with limited sample sizes or resources. To our knowledge, there is one previous study that uses a protocol developed for extraction of both CYP2E1- and GST-mediated metabolites (Kim, Collins, et al. 2009). We expanded on this method by evaluating additional metabolites, including monochloroacetic acid (MCA), trichloroethanol (TCEOH), trichloroethanol glucuronide (TCEOH-gluc), and N-acetyl-S-(1,2-dichlorovinyl)-L-cysteine (NAcDCVC), to provide greater coverage. We optimized a TCE metabolite-centered extraction protocol to detect CYP2E1- and GST-mediated metabolites using liquid chromatography mass spectrometry (LC-MS) and assessed their detection on an Orbitrap IQ-X system using a variety of columns, ion source settings, and chromatographic conditions. We then verified the extraction protocol efficiency across mouse brain, liver, and plasma matrices and assessed variability in compound detection across matrices.

To determine whether TCE metabolites can mediate PD-related neurotoxicity, we quantified TCE metabolite levels in mouse plasma, liver tissue, and brain tissue after a human-relevant inhalation exposure. Additionally, we assessed changes in global metabolism following TCE exposure in brain tissue. Our results improve our ability to detect TCE metabolites and identify likely TCE metabolite candidates that may contribute to TCE-mediated neurodegeneration.

## Materials and Methods

### Chemicals

Chemical standards for MCA, dichloroacetic acid (DCA), TCA, TCEOH, rafoxanide, and spiroxamine were purchased from Sigma-Aldrich (St. Louis, MO). TCEOH-gluc and NAcDCVC were purchased from LGC Standards (Wesel, Germany). S-(1,2-dichlorovinyl)-L-cysteine was purchased from MedChemExpress (Monmouth Junction, NJ). S-(1,2-dichlorovinyl)glutathione (DCVG) was purchased from ChemScene (Monmouth Junction, NJ). Internal standards [13C6]-D-glucose, [15N]-indole, [2-15N]-L-lysine, [15N]-choline, [13C5]-L-glutamic acid, [13C7]-benzoic acid, [15N]-L-tyrosine, uracil (1,3-15N2), [3,3-13C2]-cystine, [trimethyl- 13C3]-caffeine and [13C5]-L-glutamine were purchased from Cambridge Isotope Laboratories (Andover, MA). HPLC-grade formic acid, methanol, ammonium formate, and acetonitrile were purchased from Thermo Fisher (Waltham, MA). Milli-Q water, acetic acid, and ammonium acetate were purchased from Sigma-Aldrich.

### Standard Curves

Standard curves were produced using the TCE metabolite standards. For the mobile phase optimization experiments, standard curves were prepared such that the solvent used to dissolve the standards matched the starting mobile phase composition to be tested. The concentration range for the standard curves used to optimize the mobile phase composition was from 10 nM to 1 mM.

For the matrix-matched experiments, a new standard curve was prepared using unexposed mouse brain tissue, liver tissue, and human plasma at a concentration range between 10 nM to 100 µM. A neat standard curve of the same concentrations was prepared to perform matrix-matching calculations.

For the real samples, a matrix-matched standard curve was prepared. This matrix-matched standard curve followed the same extraction protocol as the real samples, except that they were spiked with known quantities of TCE metabolites upon reconstitution of the dried extracts, yielding a final concentration range between 10 nM to 100 μM. A neat standard curve with the same concentrations was prepared and run in parallel for quality control purposes.

Additional information on the preparation of these standard curves is provided in the supplementary information (SI1).

### Chromatography and Instrument Settings

Analyte separation was performed with a Waters CORTECS Premier C18 column (2.1 × 150 mm) in an Acquity I class HPLC system coupled to an Orbitrap IQ-X (Thermo Fisher Scientific, Waltham, MA, USA) equipped with a heated electrospray ionization probe. The column was maintained at 30 °C, with a flow rate of 0.4 mL/min and a sample injection volume of 10 μL. The mobile phase gradient was as follows: 0 - 1 min, 10% B; 1 - 7.5 min, 100% B; 7.5 - 10 min, 100% B; 10 - 10.1 min, 10% B; and 10.1 - 15 min, 10% B. The mobile phases used for each column and ionization mode are provided in Table 1.

**Table 1.** Mobile phase A and B for each experiment by column and ionization mode.

| Experiment | Condition | Mobile Phase A | Mobile Phase B |
| --- | --- | --- | --- |
| Mobile Phase Optimization, Matrix-Matching | C18Pos / C18 Neg | 0.1% Formic acid in water | Acetonitrile |
| Mobile Phase Optimization | C18Pos / C18 Neg | Water | Methanol |
| Mobile Phase Optimization | C18Pos / C18 Neg | 10 mM Ammonium acetate in water | 10 mM Ammonium acetate in acetonitrile |

The starting mobile phase composition was 90% A and 10% B, regardless of the ionization mode.

The parameters of the heated ESI (H-ESI) source were set as follows: sheath gas flow rate at 45 arbitrary units (au), auxiliary gas flow rate at 15 au, sweep gas flow rate at 0 au, RF Lens level at 55%, spray voltage at 2300 V for negative mode or 3500 V for positive mode, ion transfer tube temperature at 200 °C, and vaporizer temperature at 250 °C. The mass spectrometer was operated in data-dependent acquisition mode with a mass range of 50 – 1000 m/z and a maximum injection time of 100 ms, at a resolution of 120,000 in full-scan mode and 30,000 for MS2. These ion source parameters were determined using a direct infusion method, in which a mixture of all TCE metabolites at 500 nM was infused at 5 µL/min directly into the mass spectrometer, and source parameters were systematically varied to identify settings that maximized detection for most compounds. These optimized ion source parameters were used across all experiments.

### Internal Standards

Two pre-injection internal standards, spiroxamine and rafoxanide, were used to account for inter-injection variability. As neither compound ionizes well in both negative and positive ionization modes, spiroxamine was used for positive ionization, while rafoxanide was used for negative ionization. These compounds were validated for stability and ionizability using optimized chromatography and source parameters. A 10 µL volume of internal standards mix from a 1500 ppb stock was spiked into each sample after extraction, yielding a final concentration of 150 ppb. The following isotopically labeled standards were spiked into matrix-matched and real samples prior to extraction: [13C6]-D-glucose, [15N]-indole, [2-15N]-L-lysine, [15N]-choline, [13C5]-L-glutamic acid, [13C7]-benzoic acid, [15N]-L-tyrosine, uracil (1,3-15N2), [3,3-13C2]-cystine, [trimethyl- 13C3]-caffeine and [13C5]-L-glutamine. Final concentrations of these standards are provided in the supplementary information (SI2).

### Matrix-Matching Experiments

Using unexposed mouse brain tissue, mouse liver tissue, and human plasma, spiked matrix samples were prepared in triplicate, both pre- and post-extraction. Analytes of interest were spiked to final concentrations of 100 µM, 10 µM, 1 µM, 100 nM, and 10 nM.

Tissue samples were homogenized using a bead beater and HPLC-grade acetonitrile, such that there was 8 μL of acetonitrile to every 1 mg of tissue and 2.5 mg of 1.2 - 1.4 mm zirconium oxide beads to every 1 mg of tissue. 200 μL (25 mg) of homogenate was mixed with 660 μL of internal standards-fortified acetonitrile and 190 μL of 0.1% formic acid in water. For pre-extraction spiked matrix-matched calibrations, 15 uL of the TCE metabolite mix at the concentration needed to reach the desired final concentration was added. All samples were vortex mixed for 10 minutes, followed by centrifugation at 21,130 × g for 5 minutes. Following centrifugation, 350 µL of the supernatant was collected and evaporated to complete dryness under a nitrogen stream For the pre-extraction spiked samples, the residue was reconstituted in 100 µL of mobile phase, consisting of 90 µL of mobile phase A (0.1% formic acid in water) and 10 µL of mobile phase B (acetonitrile) fortified with internal standards rafoxanide and spiroxamine, prior to LC-HRMS analysis. For the post-extraction spiked samples, the residue was reconstituted in 100 µL of mobile phase, consisting of 85 µL of 0.1% formic acid in water, 5 µL of TCE metabolite mix (90:10, 0.1% formic acid in water:acetonitrile), and 10 µL of acetonitrile fortified with rafoxanide and spiroxamine.

Plasma was thawed on ice for 30 minutes prior to analyte extraction. For pre-extraction spiked samples, 50 uL of plasma was transferred to a clean Eppendorf tube and mixed with 116.25 μL of 0.1% formic acid in water + internal standard-fortified acetonitrile (90:10) and 8.75 μL of the TCE metabolite mix. For the post-extraction spiked samples, 50 μL of plasma was transferred to a clean Eppendorf tube and mixed with 125 μL of 0.1% formic acid in water + internal standard-fortified acetonitrile (90:10). In this instance, the internal standard-fortified acetonitrile and water mix included spiroxamine and rafoxanide. Each sample was vortexed for 1 minute and then equilibrated on ice for 30 minutes. Samples were then centrifuged for 20 minutes at 18,407 × g, maintained at 4 °C. For pre-extraction spiked samples, 100 μL of the resulting supernatant was transferred to clean LC-MS vials. For the post-extraction spiked samples, 95 μL of the resulting supernatant was transferred to clean LC-MS vials, and 5 μL of TCE metabolite mix (90:10, 0.1% formic acid in water:acetonitrile) was added. Additional information on generating the matrix-matched samples is provided in the supplementary information (SI3).

### Mouse Husbandry

Adult (9 - 12-month-old) male and female mice on a C57BL/6 background were bred at the University of Alabama at Birmingham (UAB) within the Animal Resource Program (ARP) by Dr. Briana De Miranda’s lab. All mouse inhalation exposures were conducted at UAB, using previously described methods (Adamson et al. 2023). Briefly, mice were provided with standard rodent chow and filtered water throughout the study. The mice were maintained at 28 °C and 56% humidity. At study termination, mice were humanely euthanized using a dose of CO_2_ as an anesthetic and rapid decapitation, followed by organ (liver and brain) and plasma collection. All animal experiments were conducted with existing approval by the University of Alabama at Birmingham Institutional Animal Care and Use Committee (IACUC). Mice were placed in a whole-body passive exposure chamber inside a chemical fume hood. Control animals were placed in an adjacent fume hood with a passive flow of HEPA-filtered room air. Mice were exposed to 100 ppm TCE for 4 hours, resulting in a time-weighted average of 45.126 ppm (Figure 1A). TCE concentrations were monitored using a photoionization detector (PID). The 100 ppm dose was chosen based on allometric scaling calculations, indicating that the 100 ppm dose in mice is equivalent to an 8 ppm dose in humans, which is below the workplace exposure limit set by many US regulatory agencies. In addition, the exposure used in this study falls well below the Occupational Safety and Health Administration’s (OSHA) 8-hour time-weighted average limit for TCE, which is 100 ppm (Occupational Safety and Health Administration [OSHA] 2021). Organs and plasma were collected at 0, 1, 2, 4, 8, and 24 hours post the end of exposure (Figure 1B).

**Figure 1.**
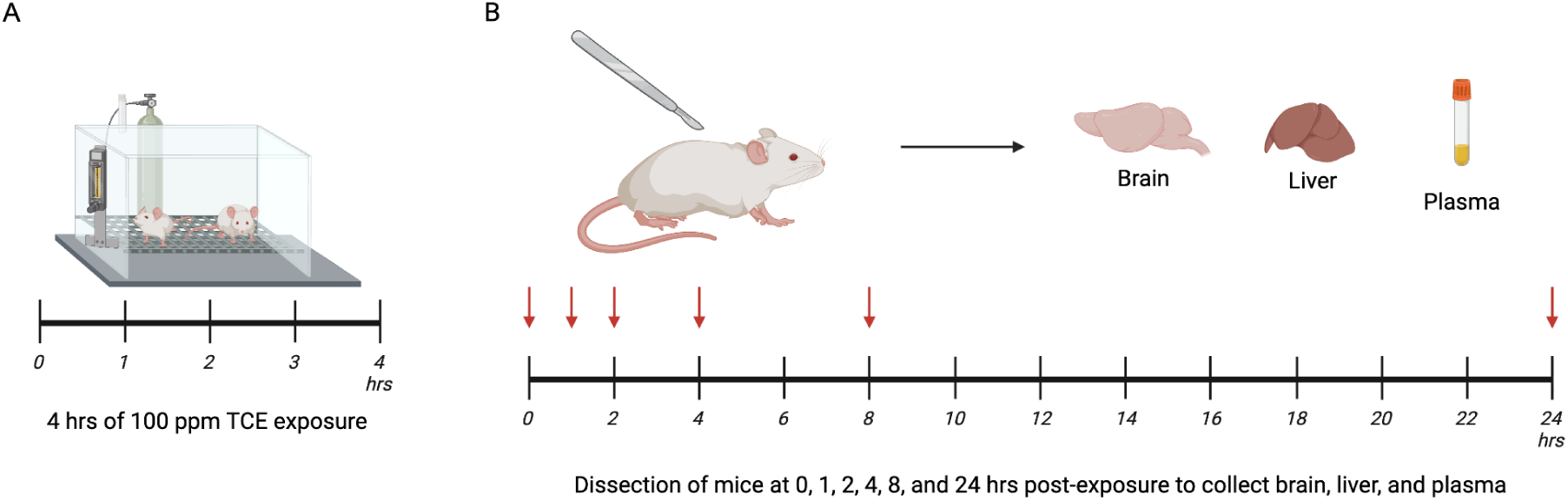
Dosing and collection scheme for mouse study. **(A)** Mouse exposure to 100 ppm TCE in a whole-body passive exposure chamber for 4 hours. **(B)** Dissection of mice at 0, 1, 2, 4, 8, and 24 hours after the end of the exposure period. Brain, liver, and plasma were collected from all animals. 5 to 6 animals per time point were dissected.

### Quantification of TCE Metabolites

Tissue levels of TCE metabolites were measured as follows. Tissue samples were homogenized using a bead beater and HPLC-grade acetonitrile, such that there was 8 μL of acetonitrile to every 1 mg of tissue. 200 μL (25 mg) of homogenate was mixed with isotopically labeled internal standard-fortified acetonitrile and water, then vortex-mixed for 10 minutes and centrifuged at 21,130 × g for 5 minutes. Following centrifugation, 350 µL of the supernatant was collected and concentrated to complete dryness under a nitrogen stream. The resulting residue was reconstituted in 100 µL of starting mobile phase, consisting of 90 µL of mobile phase A (0.1% formic acid in water) and 10 µL of mobile phase B (acetonitrile, containing pre-injection internal standards rafoxanide and spiroxamine), prior to LC-HRMS analysis.

Plasma levels of TCE metabolites were measured according to the following methods. Plasma was thawed on ice for 30 minutes before analyte extraction. 50 μL of plasma was transferred to a clean Eppendorf tube and mixed with 125 μL of a 90:10 mixture of pre-injection and isotopically labeled internal-standard-fortified acetonitrile and water containing 0.1% formic acid. Each sample was vortex-mixed for 1 minute and then equilibrated on ice for 30 minutes.

Following this equilibration period, samples were centrifuged for 20 minutes at 18,407 × g, maintained at 4 °C. 100 μL of the resulting supernatant was pipetted into clean LC-MS vials for analysis.

### HRMS-based Nontargeted Metabolomic Analysis

Extraction of samples for nontargeted analysis in mice followed a procedure similar to that used to quantify TCE metabolites, except that it employed a dual-extraction setup. This allowed for additional cleanup of the fraction partitioned for metabolomics. Due to limitations in plasma volume, this extraction protocol and analysis were performed only on liver and brain tissue.

Briefly, 200 μL of homogenate was mixed with 500 μL of acetonitrile fortified with isotopically labeled internal standards, 250 μL of water, and 500 μL of cold MTBE. The samples were vortex-mixed for 10 minutes, then centrifuged at 21,130 × g for 5 minutes. After phase separation occurred, the organic layer was removed and set aside for future lipidomic analysis.

200 μL of the aqueous layer was collected and concentrated to complete dryness under a nitrogen stream. The resulting residue was reconstituted in 100 µL of the starting mobile phase, consisting of 90 µL of mobile phase A (0.1% formic acid in water) and 10 µL of mobile phase B (acetonitrile, containing pre-injection internal standards rafoxanide and spiroxamine), prior to LC-HRMS analysis. Samples were analyzed using the same chromatography and ion source settings as noted above.

### Statistical Analysis for Mobile Phase Composition and Matrix-Matching Experiments

Using EnviPat, an isotope pattern calculator, we identified compounds of interest based on expected m/z values in either ionization mode and the matching relative abundance of chlorine isotopes (Loos et al. 2015). For mobile phase composition experiments, analytes were identified and quantified using EL-MAVEN (Agrawal et al. 2019), an open-source platform for spectral feature identification and quantification. For matrix-matching experiments, including both matrix-matched samples and neat standards, compounds were identified and quantified using Thermo Scientific Trace Finder software, which enables automated detection and peak integration of multiple compounds. Data cleaning and calculations were performed in RStudio (Version 2025.09.2+418), including normalization to internal standards and the calculation of percent recovery and matrix effects. Percent recovery, which measures the efficiency of the extraction method and how much of the analyte is successfully recovered, was calculated using the following equation:

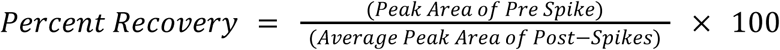

Matrix effects, which refer to interference caused by co-extracted sample components such as proteins, lipids, and salts, were calculated using the following equation:

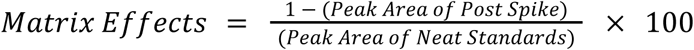

Data were plotted using the GraphPad Prism 11 statistical software package (San Diego, California).

### Statistical Analysis for Targeted Analytes

Peak areas were quantified using the Thermo TraceFinder software. Blank subtraction and data normalization to internal standards were conducted using RStudio. Matrix-matched calibration curve values were fitted using ordinary least squares (OLS) linear regression with a zero-intercept model for each compound. The resulting equations were used to predict concentrations in the samples. The following R packages were used to carry out these calculations: tidyverse, readxl, beepr, janitor, writexl, and broom (Wickham and RStudio 2023; Bååth and Dobbyn 2024; Firke et al. 2024; Ooms and McNamara 2025; Robinson et al. 2026; Wickham et al. 2026)

Statistical analyses of this data were performed using GraphPad Prism 11. Welch’s ANOVA followed by Dunnett’s T3 multiple-comparisons test, or one-way ordinary ANOVA followed by Tukey’s post hoc test, was used to compare analyte concentrations across time points.

In addition, toxicokinetic modeling was used to solve the system of ordinary differential equations that describe the model. Model parameters were estimated by minimizing a weighted least-squares cost function using nonlinear optimization. Residuals were normalized by the maximum observed concentration in each tissue. To ensure that modeling was possible, negative values were set to 0. The following packages were used to carry out these calculations: tidyverse, janitor, readxl, deSolve, pracma, ggh4x, and beepr (Wickham and RStudio 2023; Bååth and Dobbyn 2024; Firke et al. 2024; Brand 2025; Borchers 2025; Soetaert et al. 2026; Wickham et al. 2026). A three-compartment model was assumed, with the liver as the site of formation and plasma as the primary site of elimination. The following set of ordinary differential equations was used:

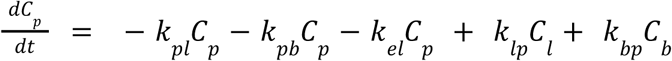

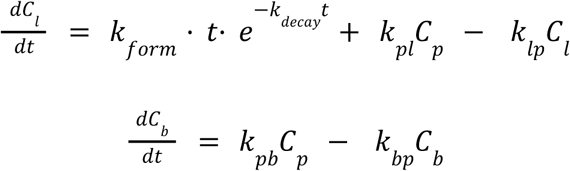

where C_p_, C_l_, and C_b_ represent concentrations in plasma, liver, and brain, respectively; k_form_ is a rate constant that represents formation of the metabolite in the liver; k_decay_ is a rate constant that represents the temporal decay of formation of the metabolite; k_pl_ and k_pb_ are rate constants for the transfer from plasma to liver and brain; k_lp_ and k_bp_ are rate constants for the transfer from liver and brain back to plasma; and k_el_ is the elimination rate constant from plasma.

Half-lives were calculated using the following equations:

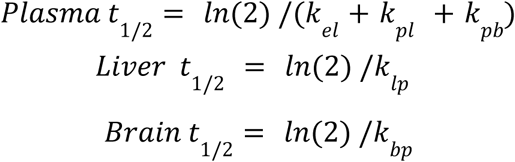

AUC was calculated from the observed data via numerical integration using the trapezoidal rule with the trapz function in the “pracma” package. An additional toxicokinetic analysis was conducted with the brain as a site of formation; details are provided in the supplementary information (SI4)

### Statistical Analysis for Nontargeted Metabolomic Analysis

Feature detection and peak integration were done using MZmine (Version 4.8.0). Feature lists were initially filtered in MZmine. The minimum number of detections in blanks was set to 3 of 5, and the quantification area was used for comparison. Features with a fold-change increase of ≥300% relative to blanks were retained. The minimum feature height was set to 1 x 10^4^.

Additional information on MZmine inputs can be found in the supplementary information (SI5). Filtered feature lists were exported for statistical testing in RStudio. To ensure that the retained features were specific to that particular experimental group, features not detected in at least 50% of the mice in that group were excluded. The data were normalized using the internal standards rafoxanide for negative ionization mode and spiroxamine for positive ionization mode. Missing values were imputed using half the minimum observed value (LOD/2) for missing or zero values, and all values were log2-transformed.

Data were mean-centered and scaled prior to principal component analysis (PCA), which was used to assess overall sample clustering using an unsupervised method. Partial least squares discriminant analysis (PLS-DA) was performed on log2-transformed data as a supervised method to evaluate sample clustering. Differential abundance analysis was performed using pairwise unpaired Welch’s t-tests. P-values were adjusted for multiple comparisons using the Benjamini-Hochberg false discovery rate method. Differential abundance analysis was performed separately for each mode within each matrix. This was conducted using the following R packages: tidyverse, janitor, mixOmics, ggrepel, beepr, and ggprism (Wickham and RStudio 2023; Bååth and Dobbyn 2024; Firke et al. 2024; Dawson 2025; Slowikowski et al. 2026; mixOmics).

The timepoint comparison with the greatest number of significant differences, control vs. 4 hr post-exposure, was used for downstream analyses. For this analysis, m/z, retention time, p-value, and t-statistic values were retained from the differential abundance analysis results.

Results from both ionization modes were aggregated for pathway enrichment analysis to improve metabolome coverage. Pathway analysis was performed using the functional analysis module of the MetaboAnalyst 6.0 web server (https://www.metaboanalyst.ca/). The Mummichog algorithm was used with BioCyc and KEGG annotation libraries for *Mus musculus* (house mouse).

Additional information on MetaboAnalysis inputs and the results for the secondary analysis using liver tissue is included in the supplementary information (SI6 and SI7).

## Results

We developed a method to maximize detection of both oxidative (TCA, DCA, MCA, TCEOH, and TCEOH-gluc) and (DCVC, DCVG, and NAcDCVC) glutathione-conjugated metabolites via a single extraction protocol, expanding upon previous publications that report detection and quantification of TCE metabolites using multiple methods to detect both types of metabolites (Bernauer et al. 1996; Ketcha et al. 1996; Brashear et al. 1997; Lash et al. 1999; Merdink et al. 2008; Venkatratnam et al. 2018; Luo, Hsieh, et al. 2018b). To do this, we optimized the ion source settings via direct infusion and then optimized the mobile phase composition to improve compound separation and retention. Then, we calculated the percent recovery and matrix effects for our extraction method. Finally, we applied our method to real biological samples from the TCE inhalation mouse cohort and assessed them using targeted and nontargeted analyses.

### Mobile Phase Optimization

We tested three mobile phase combinations: acidic (0.1% formic acid in water and acetonitrile), neutral (methanol and water), and basic (10 mM ammonium acetate in water and 10 mM ammonium acetate in acetonitrile). TCE metabolite standards were dissolved in the starting mobile phases for each tested mobile-phase combination and run on the instrument using the optimized ion source settings on a C18 column in both negative and positive ion modes. First, we assessed the compounds detected only in negative ionization mode (Figure 2). We found that, on average, detection sensitivity was improved under acidic mobile phase conditions compared with basic and neutral conditions, as indicated by increased peak areas of the analytes.

**Figure 2.**
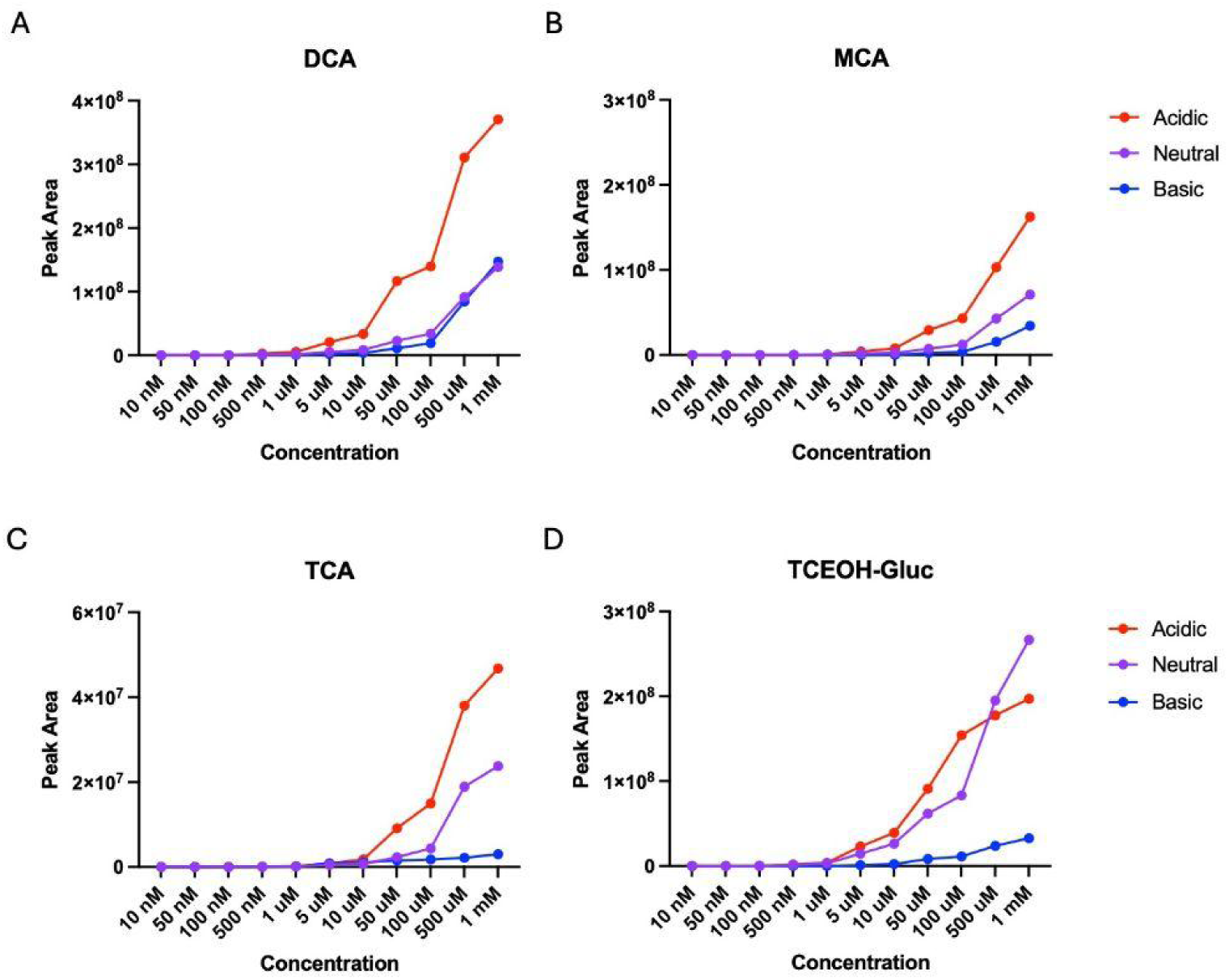
Peak areas of TCE metabolites detected in negative ionization mode using acidic, neutral, and basic mobile phase compositions. Detection of (A) DCA, (B) MCA, (C) TCA, and (D) TCEOH-Gluc from 10 nM to 1 mM across all mobile phase conditions. The starting mobile phase composition for all conditions was 90% A and 10% B. Acidic mobile phase denotes 0.1% formic acid in water (mobile phase A) and acetonitrile (mobile phase B). Neutral mobile phase denotes water (mobile phase A) and methanol (mobile phase B). Basic mobile phase denotes 10 mM ammonium acetate in water (mobile phase A) and 10 mM ammonium acetate in acetonitrile (mobile phase B). N = 1 per concentration per mobile phase composition.

Next, we assessed the compounds that could be detected in both negative and positive ionization modes (Figure 3). In general, we observed that although the compounds could be detected in both ionization modes, they were more sensitive in positive ionization mode. In addition, we observed that detection sensitivity was higher under acidic mobile-phase conditions than under basic or neutral conditions for all compounds in positive ionization mode.

**Figure 3.**
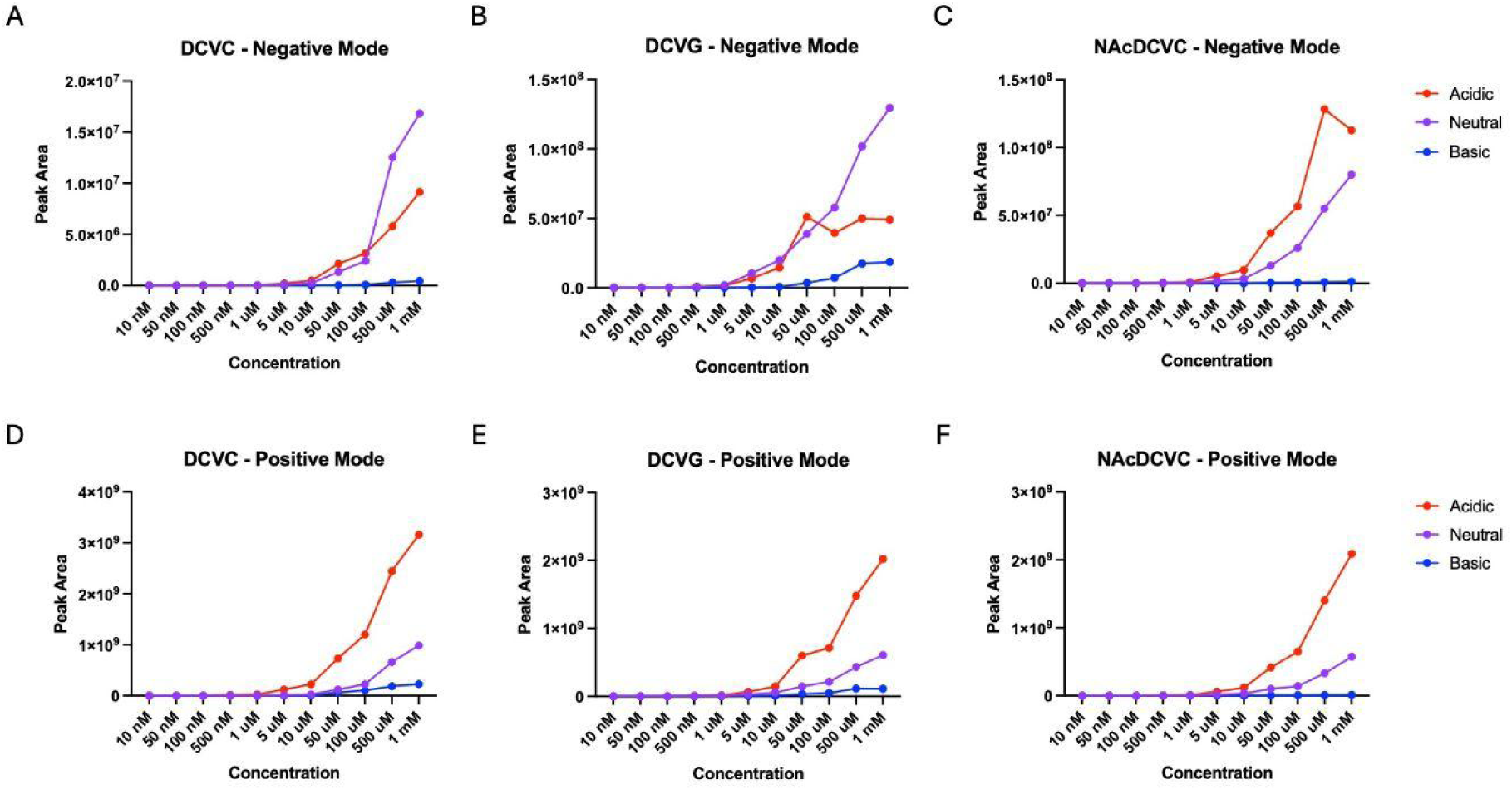
Peak areas of TCE metabolites detected in negative and positive ionization mode using acidic, neutral, and basic mobile phase compositions. Detection of (A) DCVC in negative mode, (B) DCVG in negative mode, (C) NAcDCVC in negative mode, (D) DCVC in positive mode, (E) DCVG in positive mode, and (C) NAcDCVC in positive mode across all mobile phase conditions. The starting mobile phase composition for all conditions was 90% A and 10% B. Acidic mobile phase denotes 0.1% formic acid in water (mobile phase A) and acetonitrile (mobile phase B). Neutral mobile phase denotes water (mobile phase A) and methanol (mobile phase B). Basic mobile phase denotes 10 mM ammonium acetate in water (mobile phase A) and 10 mM ammonium acetate in acetonitrile (mobile phase B). N = 1 per concentration per mobile phase composition.

### Percent Recovery and Matrix Effects

After column chromatography and mass spectrometer settings were optimized, the detection of these compounds in the matrix was assessed using unexposed mouse brain, mouse liver, and human plasma. With this in mind, we conducted a series of matrix-matched experiments in which we spiked a matrix with known concentrations of the compounds and then assessed percent recovery and matrix effects. We selected 10 nM, 100 nM, 1 µM, 10 µM, and 100 µM as the concentrations of interest for matrix matching. With this in mind, first we assessed the percent recovery by identifying and quantifying the compounds (Figure 4). We found that in the liver, most compounds had a percent recovery of at least 50%, with a few exceptions. In many cases, compounds detected in both negative and positive ionization modes had higher percent recovery in positive ionization mode (Figure 4A). In addition, we generally found that percent recovery increased with concentration, except for TCA, which declined sharply at the highest concentration. The brain tissue showed very similar trends, and we also noted that the compounds we could detect in both appeared to be better in positive ionization mode (Figure 3B). Finally, in plasma, we found that percent recoveries were generally similar, though a few concentrations showed extreme values well above 100% (Figure 4C). In plasma, the percent recoveries on the lower end of the concentration curve were not detected, so overall there was poorer detection of the compounds in plasma as compared to the other matrices at the lower end. Additionally, several plasma concentrations showed percent recoveries well above 100%, suggesting possible ion enhancement.

**Figure 4.**
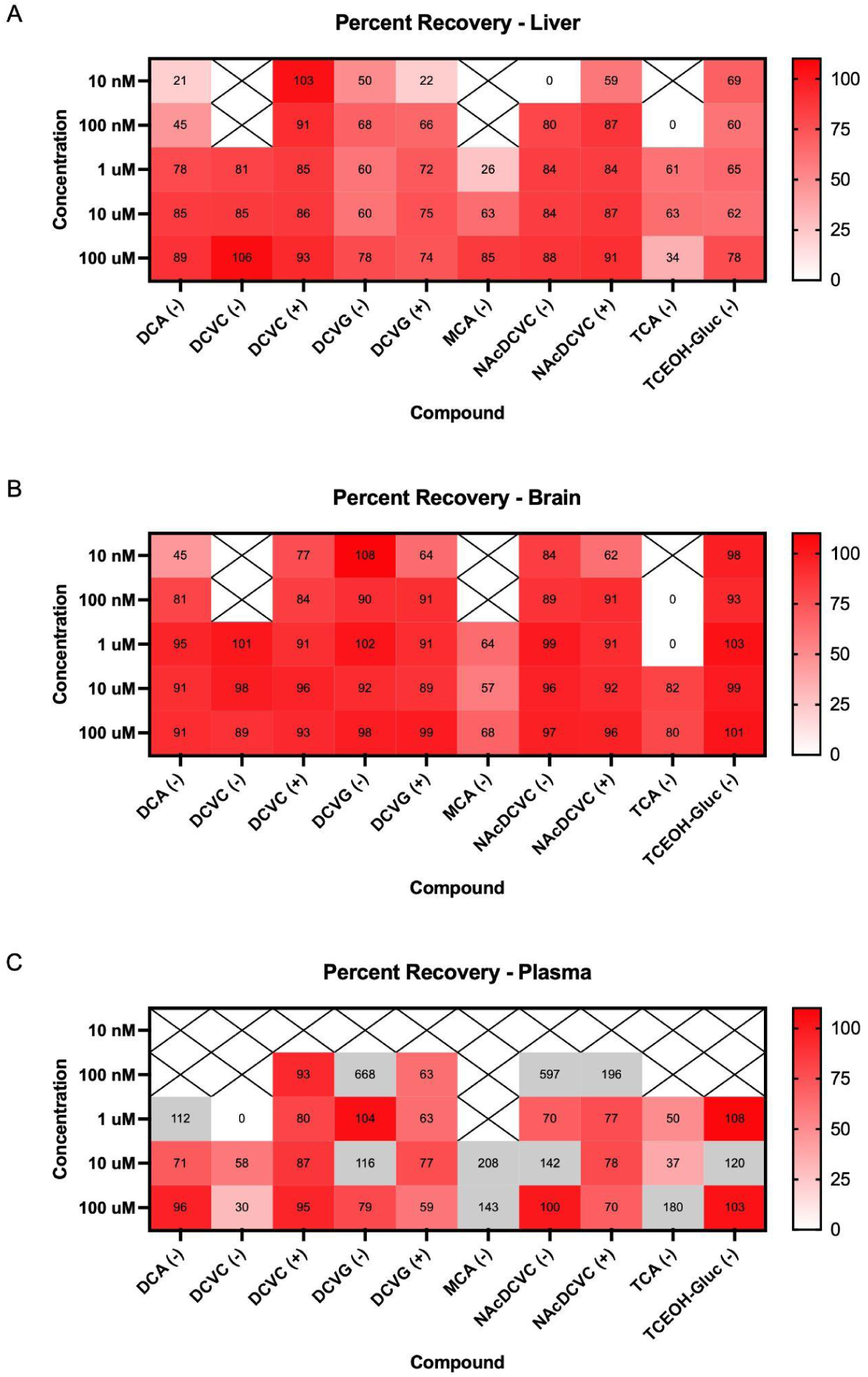
Percent recoveries of TCE metabolites across matrices. Percent recoveries in (A) liver, (B) brain, and (C) plasma. The (+) or (-) next to each analyte denotes that it was measured in positive or negative mode, respectively. For consistency, the same color gradient range was applied to each graph. Values that fall outside of this range are colored in gray. A white box with an ’x’ indicates that the compound was not detected at that concentration. N = 3 per concentration.

In addition to percent recovery, we assessed matrix effects for these compounds (Figure 5). In the liver, we found that several compounds exhibited signal suppression or matrix effect values greater than 0% across most or all concentrations, including DCVC, DCVG, NAcDCVC, MCA, and TCEOH-Gluc (all detected in negative ion mode) (Figure 5A). Otherwise, we observed signal enhancement for most of the other compounds. We observed a similar trend in the brain, although for TCEOH-Gluc and NAcDCVC (detected in negative ion mode), we found substantially enhanced signals across concentrations for both compounds (Figure 5B). There is also signal suppression for TCA. For plasma, most compounds across all concentrations exhibit signal suppression, except for TCA, which shows substantially greater signal enhancement than any other compound or matrix (Figure 5C).

**Figure 5.**
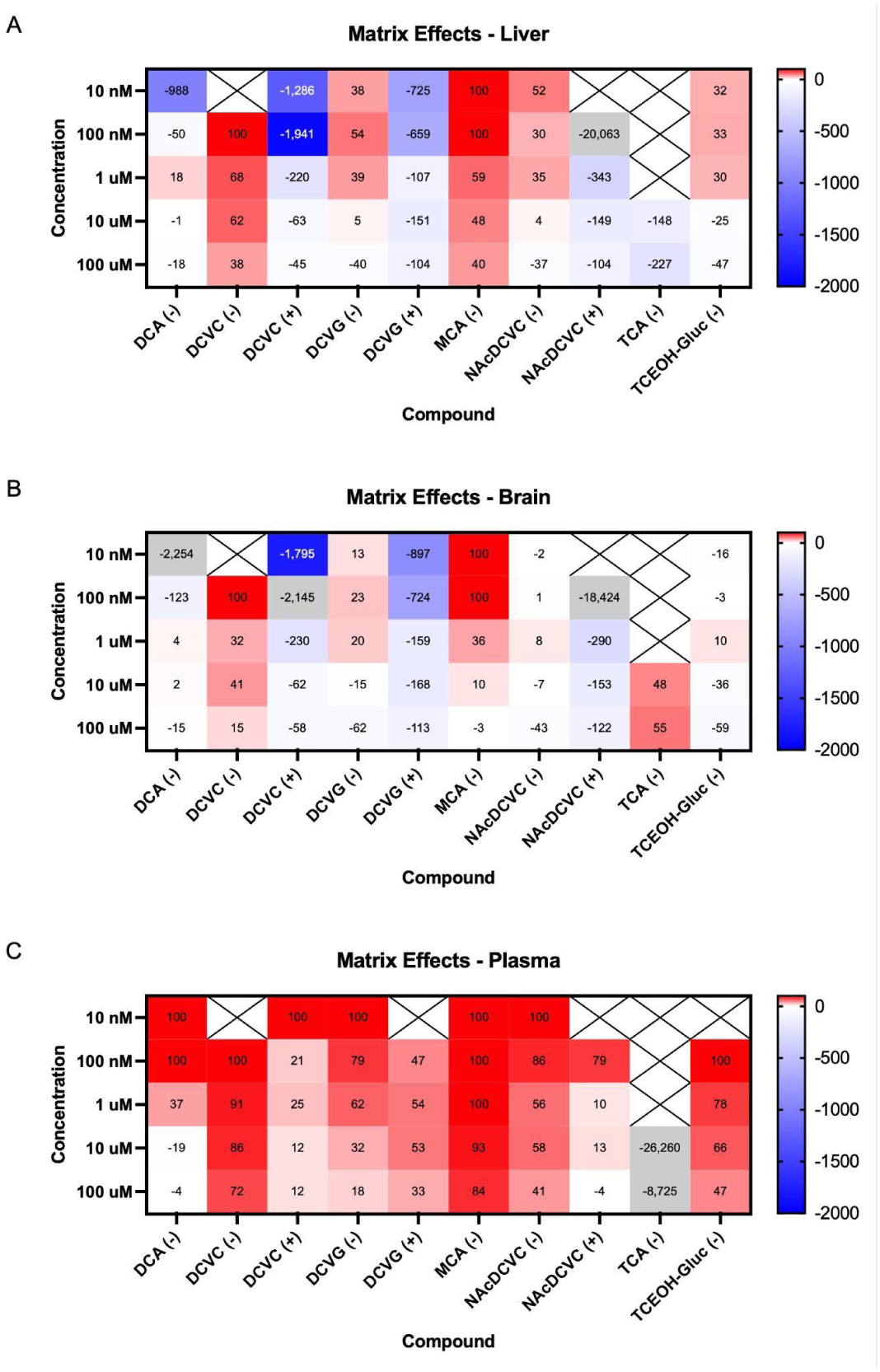
Matrix effects on TCE metabolites across matrices. Matrix effects in (A) liver, (B) brain, and (C) plasma. The (+) or (-) next to each analyte denotes that it was measured in positive or negative mode, respectively. Negative percentages indicate signal enhancement, while positive percentages indicate signal suppression. For consistency, the same color gradient range was applied to each graph. Values that fall outside of this range are colored in gray. A white box with an ’x’ indicates that the compound was not detected at that concentration. N = 3 per concentration.

### Quantification of TCE Metabolites

Of the TCE metabolites that can be detected by our method, levels of TCE metabolites DCA, TCA, and TCEOH-gluc were quantified in mouse brain, liver, and plasma after exposure to TCE across multiple time points. TCE metabolite levels were quantified over time in the brain, liver, and plasma after 4 hours of TCE exposure, with matrix-matched calibration curves.

Concentrations are reported as mean ± standard deviation (SD). DCA was detected at low levels throughout the 24-hour assessment period (Figure 6A). DCA levels peaked at 1 and 4 hours post-exposure across all matrices. Plasma levels were the highest of all matrices at 1 hour, with a peak concentration of 0.21 ± 0.17 μM. Brain levels were the highest of all matrices at 4 hours, with a peak concentration of 0.34 ± 0.14 μM. TCA was detected at high quantities over the 24-hour assessment period (Figure 6B). TCA was detected at the highest concentrations in plasma, peaking at 1 hour post-exposure at 190.10 ± 140.30 μM. TCA peaked at 8 hours post-exposure in both the brain and the liver, with peak concentrations of 14.00 ± 3.50 μM and 37.37 ± 16.96 μM, respectively. TCEOH-Gluc was detected at high quantities over the 24-hour assessment period in both liver and plasma, and at low levels in the brain (Figure 6C).

**Figure 6.**
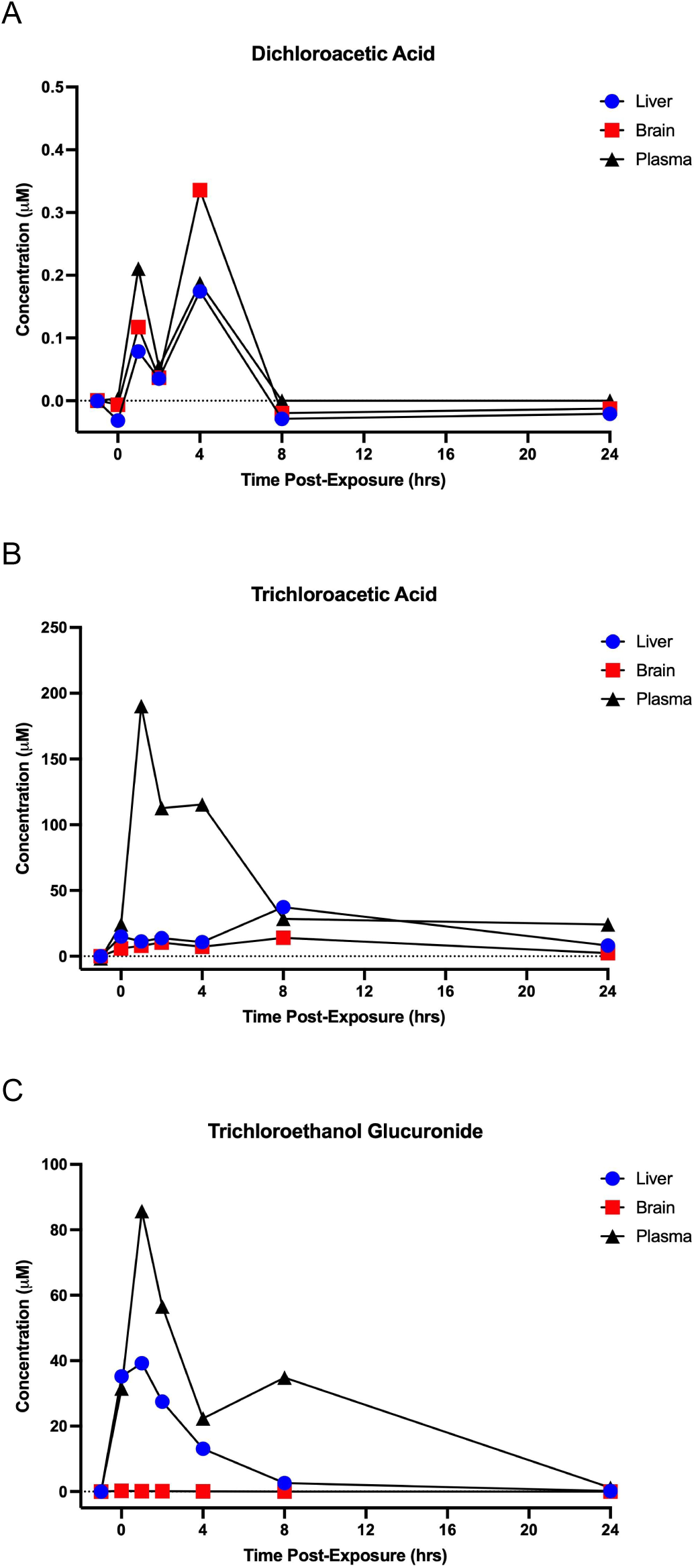
Mean values of concentrations of DCA, TCA, and TCEOH-Gluc in liver, brain, and plasma over time. Concentrations over time in liver, brain, and plasma samples for (A) DCA, (B) TCA, and (C) TCEOH-gluc. Across these graphs, values at -1 on the x-axis represent the concentrations of the analytes in the unexposed group. Liver samples are denoted using blue circles, brain samples are denoted using red squares, and plasma samples are denoted using black triangles. N = 5 - 6 mice per time point.

TCEOH-Gluc was detected at the highest concentrations in plasma, peaking at 1 hour post-exposure at 85.74 ± 39.65 μM. TCEOH-Gluc peaked in the liver at 1 hour post-exposure, with a peak concentration of 39.25 ± 4.75 μM. Although TCEOH-Gluc was detected at lower levels in the brain than in the liver and plasma, it followed a similar trend, peaking at 0 hours post-exposure at 0.21 ± 0.15 μM (Figure 7H).

**Figure 7.**
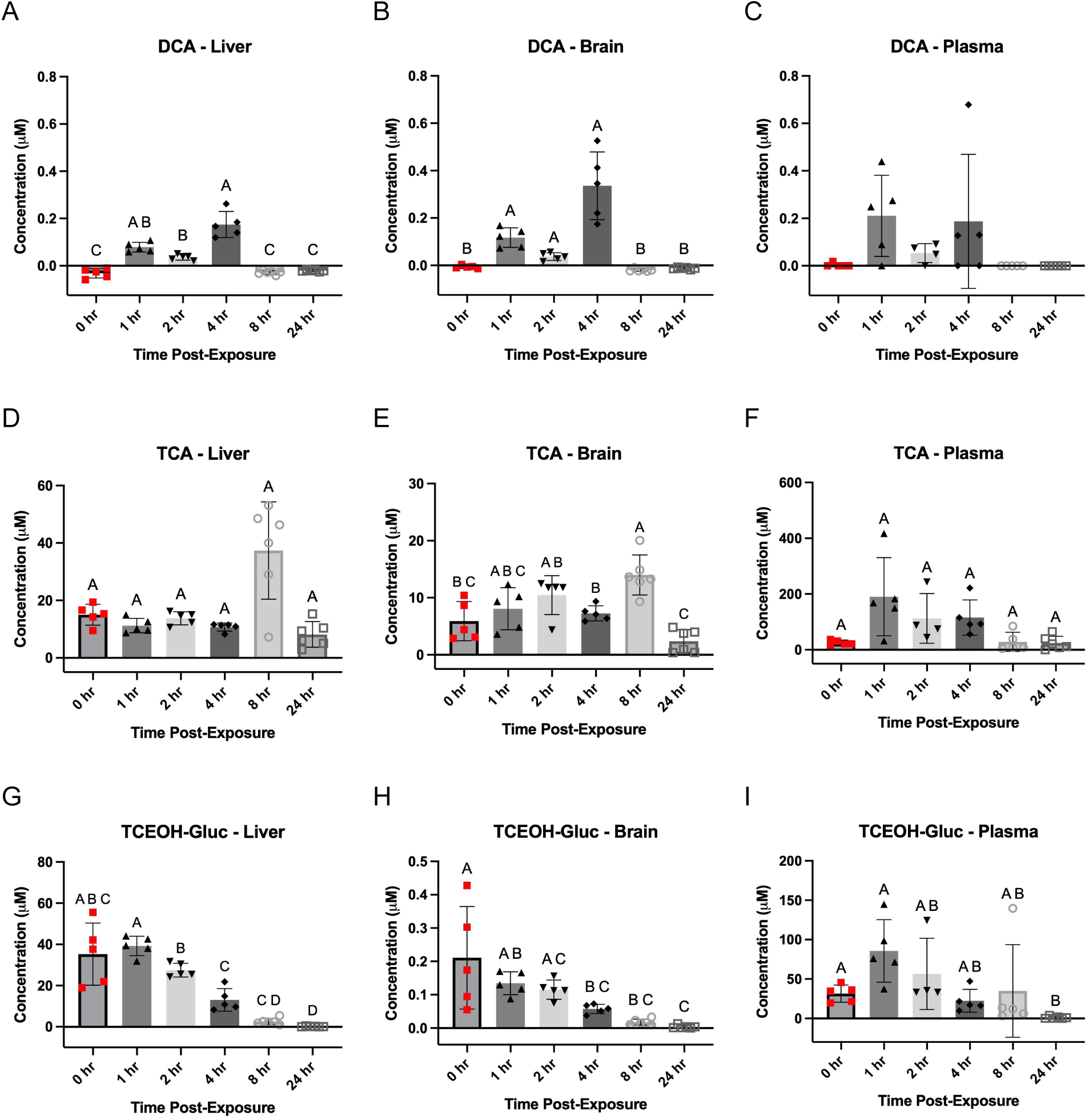
Mean and individual values of concentrations of DCA, TCA, and TCEOH-Gluc in liver, brain, and plasma over time. Error bars represent ± SD. Bars that share a letter are not significantly different from one another, using Dunnett’s T3 (p < 0.05) for Welch’s ANOVA follow-up and Tukey’s post hoc test (p < 0.05) for ordinary one-way ANOVA follow-up. **(A)** DCA in liver. Welch’s ANOVA, W(5, 10.66) = 46.82, p < 0.001. **(B)** DCA in brain. Welch’s ANOVA, W(5, 11.05) = 22.16, p < 0.001. **(C)** DCA in plasma. One-way ANOVA, F(5, 24) = 2.660, p = 0.048; no pairwise comparisons were significant after correction. **(D)** TCA in liver. Welch’s ANOVA, W(5, 11.80) = 4.636, p = 0.014; no pairwise comparisons were significant after Dunnett’s T3 correction. **(E)** TCA in brain. Welch’s ANOVA, W(5, 11.61) = 9.978, p = 0.001. **(F)** TCA in plasma. Welch’s ANOVA, W(5, 9.815) = 3.248, p = 0.054; no pairwise comparisons were significant after correction. **(G)** TCEOH-Gluc in liver. Welch’s ANOVA, W(5, 9.777) = 116.9, p < 0.001. **(H)** TCEOH-Gluc in brain. Welch’s ANOVA, W(5, 10.61) = 33.46, p < 0.001. **(I)** TCEOH-Gluc in plasma. Welch’s ANOVA, W(5, 8.833) = 11.99, p = .001. N = 4 - 6 mice per time point.

To evaluate whether these trends were statistically significant, time-dependent differences for each metabolite in each matrix were tested individually (Figure 7). As these data were blank-subtracted using values from the unexposed mouse group, the unexposed group was omitted from statistical testing. For DCA, liver and brain concentrations showed significant pairwise differences between early post-exposure timepoints (1, 2, and 4 hr) and later timepoints where concentrations were near 0 μM, indicating rapid formation and clearance of DCA in both matrices (Figure 7A and 7B). Although plasma shows a similar trend and indicates a significant difference across groups after an ordinary one-way ANOVA (p = 0.0475), the spread of the values is such that no pairwise differences are statistically significant between any of the time points (Figure 7C). The 8- and 24-hr group values were both 0 μM because DCA was not detected in any samples. The formation of DCA in such low quantities across all matrices is consistent with it being a minor metabolite of TCE, downstream of TCA. TCA can be readily excreted, making DCA formation less likely.

For TCA, brain concentrations increase over time up to 8 hr post-exposure, then significantly drop at 24 hr post-exposure (Figure 7E). Concentrations at 8 hr differed significantly from 0 and 24 hr; concentrations at 4 hr differed significantly from 8 and 24 hr; and concentrations at 2 hr differed significantly from 24 hr. The presence of TCA at the 0 hr timepoint, along with its increase up to 8 hr, suggests that TCA is formed rapidly after the start of exposure and continues to form well after the end of exposure. Its rapid formation is consistent with prior literature reporting TCA as a primary metabolite of TCE (Bernauer et al. 1996; Lash et al. 2000; Luo, Hsieh, et al. 2018b). Continued detection of TCA at the 24 hr timepoint is consistent with previously reported information on TCA’s long biological half-life. Liver concentrations of TCA remained stable across timepoints until 8 hr, after which the TCA concentration peaked before declining at 24 hr. Plasma TCA concentrations increased rapidly between 0 and 1 hr, peaking at 1 hr and then declining over time (Figure 7F). Welch’s ANOVA detected a near-significant (0.05 < *p* < 0.1) difference across timepoints (*p* = 0.0544). The near-significant p-value suggests that TCA may exhibit subtle time-dependent changes in plasma as compared to the other matrices.

For TCEOH-Gluc, liver and plasma concentrations were initially high, peaking at 1 hr, and then declined steadily until the 24 hr timepoint (Figure 7G and 7I). In the liver, concentrations at 1 hr differed significantly from 2, 4, 8, and 24 hr; concentrations at 2 hr differed significantly from 4, 8, and 24 hr; and concentrations at 0 and 4 hr differed significantly from 24 hr. Due to variability in the data, significant pairwise differences were observed in plasma between 0 and 24 hr and between 1 and 24 hr. Brain concentrations of TCEOH-Gluc followed a similar trend, peaking at 0 hr and steadily declining through 24 hr (Figure 7H).

Concentrations at 1 and 4 hr differed significantly from 8 and 24 hr; concentrations at 2 hr differed significantly from 8 and 24 hr; and the concentration at 1 hr differed significantly from 4 hr. These data suggest that TCEOH-Gluc is formed rapidly after the start of exposure but is almost completely cleared after 24 hours across all matrices. Its rapid formation and clearance across matrices are consistent with prior literature identifying TCEOH-Gluc as the main secondary metabolite of TCEOH, a primary metabolite of TCE (Stenner et al. 1997; Lash et al. 2000).

### Toxicokinetic Modeling for TCE Metabolites

Toxicokinetic modeling was performed to better understand metabolite formation and excretion (Figure 8). A series of starting parameters was tested to find the best model fit, illustrated in Figure 8. A sensitivity analysis was also performed to account for potential on-site formation of the metabolites in the brain, though this did not improve model performance for these metabolites; details are provided in the supplementary information (SI4).

**Figure 8.**
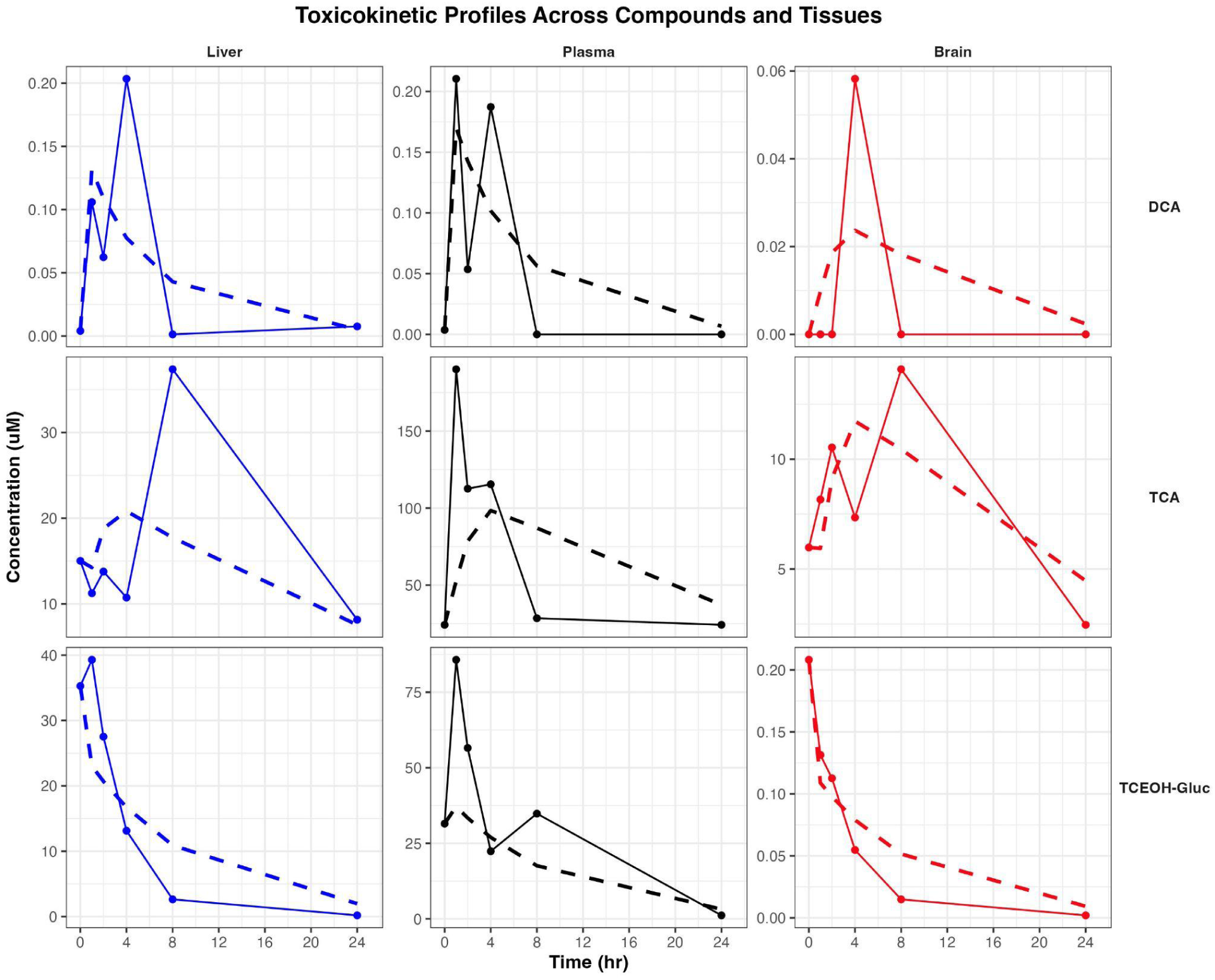
Observed and model-predicted toxicokinetic (TK) profiles across matrices. Points and solid lines represent the observed concentration-time data, while the dashed lines represent the model-predicted concentrations.

**Figure 9.**
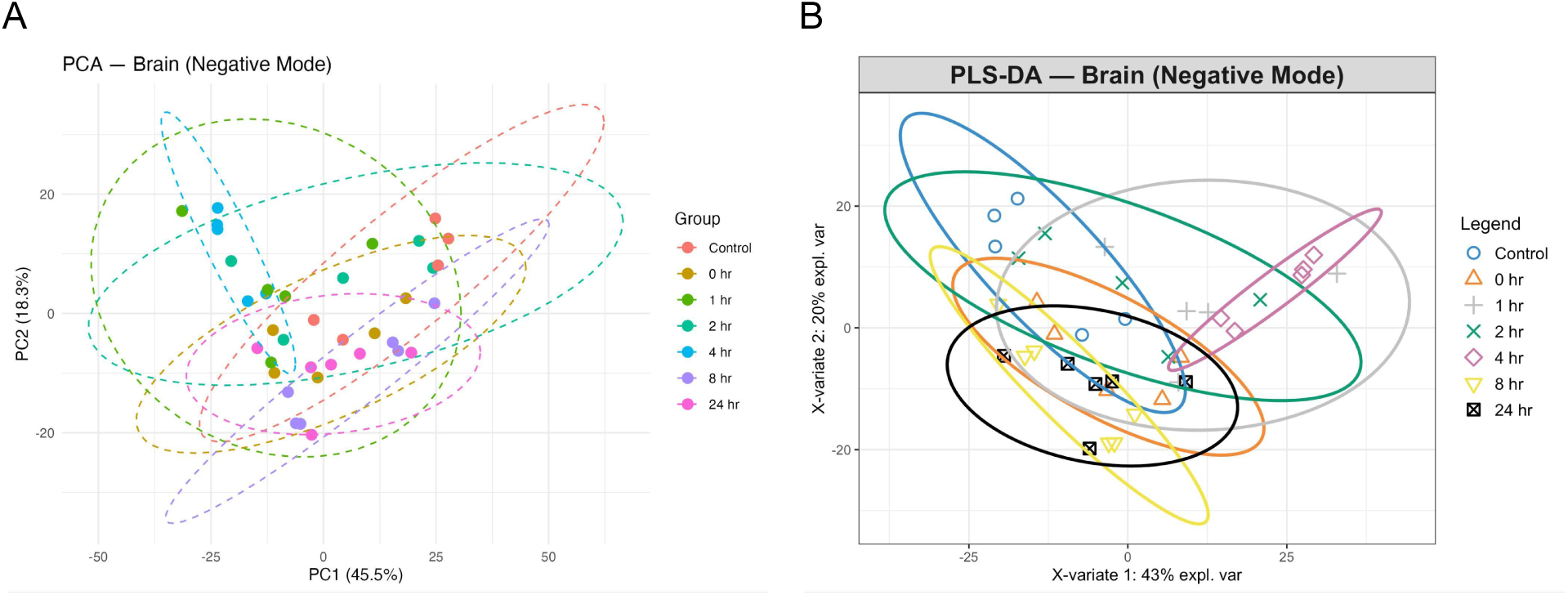
PCA and PLS-DA for all groups in negative ionization mode. **(A)** PCA and **(B)** PLS-DA. Positive ionization mode PCA and PLS-DA graphs are omitted as there was no apparent difference between them from the negative ionization mode graphs.

The following rate constants were estimated from the system of ODEs using the best-fitting model (Table 2). DCA had the highest rate constant for elimination, consistent with its relatively rapid elimination from plasma after formation. TCA had the highest rate constant for formation and the lowest for elimination, indicating that it is rapidly formed and has slower clearance than the other metabolites. TCEOH-Gluc had the highest decay rate constant, suggesting that its formation is more transient than that of the other metabolites.

**Table 2.** Estimated rate constants for DCA, TCA, and TCEOH-Gluc from the three-compartment model.

| Compound | $k_{pl}$ | $k_{pb}$ | $k_{lp}$ | $k_{bp}$ | $k_{el}$ | $k_{form}$ | $k_{decay}$ |
| --- | --- | --- | --- | --- | --- | --- | --- |
| DCA | 7.265 | 0.093 | 9.684 | 0.394 | 0.280 | 16.097 | 6.877 |
| TCA | 2.023 | 1.195 | 10.000 | 9.996 | 0.071 | 113.533 | 0.982 |
| TCEOH-Gluc | 6.131 | 0.022 | 10.000 | 7.531 | 0.173 | 0.063 | 9.999 |

Half-lives, AUCs, and AUC ratios were calculated for each metabolite within each matrix (Table 3). Half-lives were generally the highest in plasma, except for DCA, where the brain had the highest half-life. This should be considered cautiously, as the DCA in the brain model had only one nonzero time point, resulting in a somewhat poor model fit. TCA, which has the highest reported half-life in mice, continued to have the highest plasma half-life (Kim, Kim, et al. 2009). The AUC values indicate that exposure to TCA and TCEOH-Gluc was greater across all matrices as compared to DCA. The AUC ratios indicate greater exposure to these compounds in plasma than in the liver or brain. Model fit parameters were relatively good for these models, especially for TCEOH-Gluc. Model performance for TCA in plasma was poor, as indicated by the negative pseudo-R^2^ value.

**Table 3.** Toxicokinetic parameters and model performance metrics for DCA, TCA, and TCEOH-Gluc across matrices.

| Compound | Tissue | Half-life (hr) | AUC | AUC ratio vs. plasma | RMSE | NRMSE | Pseudo R<sup>2</sup> |
| --- | --- | --- | --- | --- | --- | --- | --- |
| DCA | Plasma | 0.091 | 0.854 | 1.000 | 0.058 | 0.275 | 0.578 |
|  | Liver | 0.072 | 0.886 | 1.036 | 0.058 | 0.287 | 0.360 |
|  | Brain | 1.758 | 0.175 | 0.205 | 0.018 | 0.311 | 0.302 |
| TCA | Plasma | 0.211 | 1,196.093 | 1.000 | 63.217 | 0.333 | - 0.030 |
|  | Liver | 0.069 | 510.611 | 0.427 | 9.325 | 0.250 | 0.092 |
|  | Brain | 0.069 | 209.601 | 0.175 | 2.691 | 0.191 | 0.448 |
| TCEOH-Gluc | Plasma | 0.110 | 611.086 | 1.000 | 23.164 | 0.270 | 0.247 |
|  | Liver | 0.069 | 165.417 | 0.271 | 8.117 | 0.207 | 0.718 |
|  | Brain | 0.092 | 0.734 | 0.001 | 0.021 | 0.102 | 0.912 |

### Nontargeted Analysis of Global Metabolism

In addition to quantifying TCE metabolite levels, an analysis was conducted to assess changes in global metabolism in liver and brain tissue. Plasma was not included due to limited sample volume. Given that the relevance of global metabolic changes in the brain post-TCE exposure far outweighs that of the liver, only the brain results are reported. Liver results can be found in the supplementary information (SI7).

Results of the PCA and PLS-DA indicate weak separation among the groups (Figure 3.5).

However, the PLS-DA indicates that the separation of the 4 hr group from the control is more pronounced than at other time points, which was one factor that motivated the selection of the control vs. 4 hr contrast for downstream analyses.

Following the PCA and PLS-DA analysis, multiple comparisons were performed by comparing each time point to all other time points using Welch’s unpaired t-test and the Benjamini-Hochberg false discovery rate method. This analysis was initially conducted separately for negative and positive ionization modes. To specifically examine differences between unexposed controls and exposed groups, volcano plots were generated to visualize significant features across time points relative to the controls (Figures 10 and 11). The 4 hr time point had the greatest number of statistically significant features compared to the control group in both positive and negative ionization modes.

**Figure 10.**
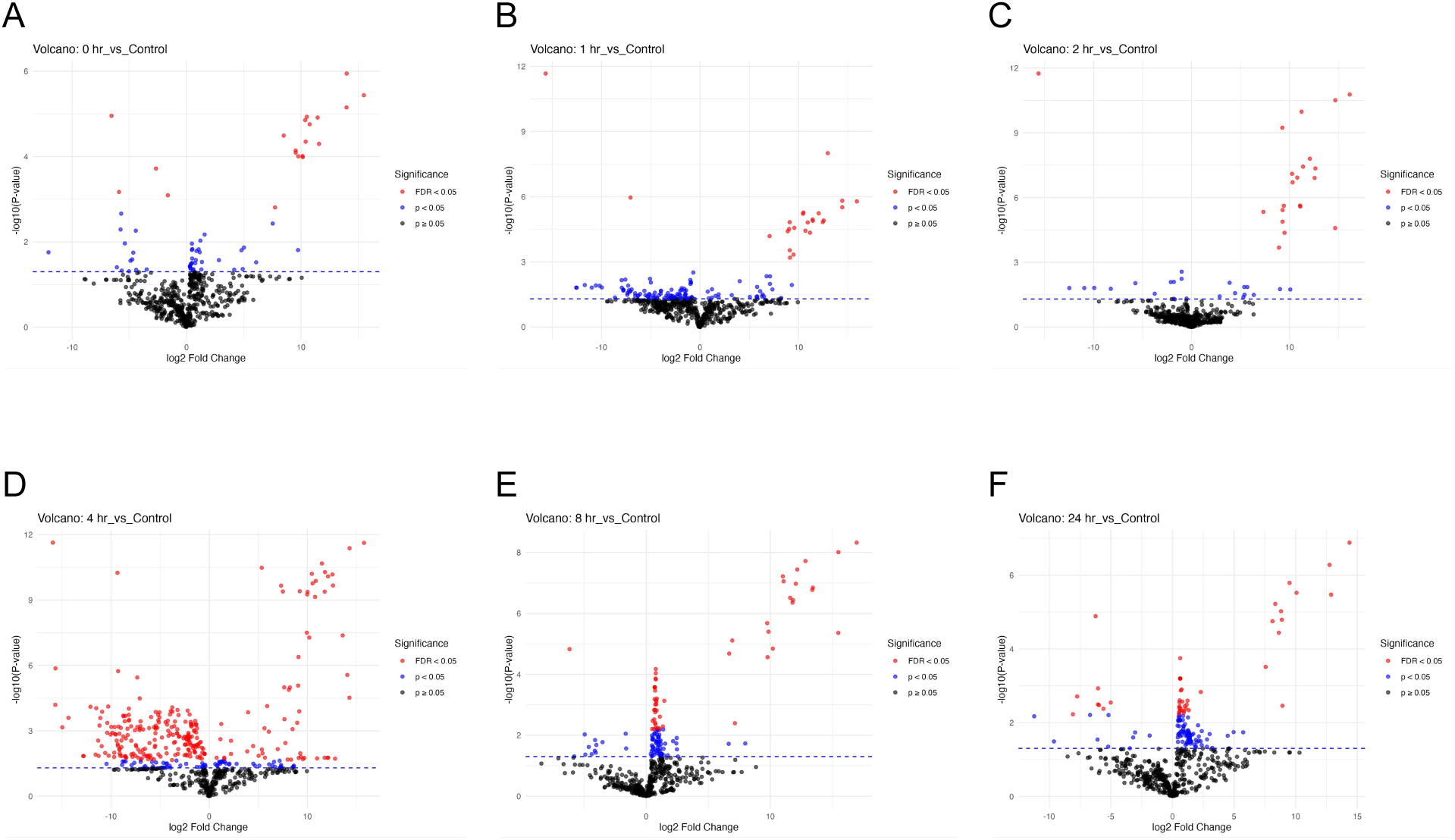
Comparison of the abundances of features in the unexposed control group to the experimental groups in negative ionization mode. Differential abundance of control vs. **(A)** 0 hr post-exposure group, **(B)** 1 hr post-exposure group, **(C)** 2 hr post-exposure group, **(D)** 4 hr post-exposure group, **(E)** 8 hr post-exposure group, and **(F)** 24 hr post-exposure group. N = 5 - 6 mice per group.

**Figure 11.**
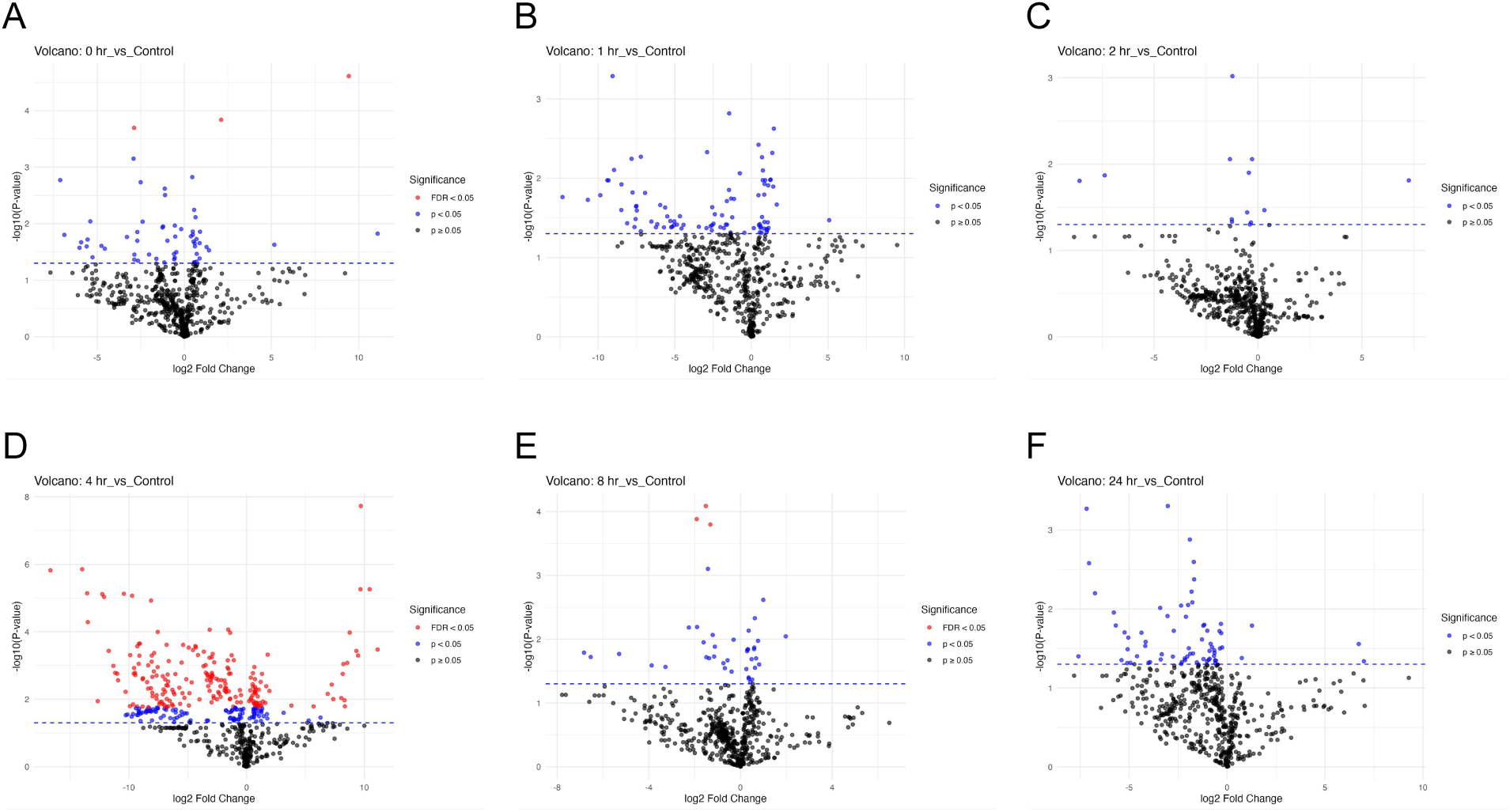
Comparison of the abundances of features in the unexposed control group to the experimental groups in positive ionization mode. Differential abundance of control vs. **(A)** 0 hr post-exposure group, **(B)** 1 hr post-exposure group, **(C)** 2 hr post-exposure group, **(D)** 4 hr post-exposure group, **(E)** 8 hr post-exposure group, and **(F)** 24 hr post-exposure group. N = 5 - 6 mice per group.

Since the 4 hrs post-exposure period showed the greatest change in global metabolism compared with unexposed controls, functional analysis was performed on this comparison using the MetaboAnalyst 6.0 web server. This analysis was performed using the mixed-mode setting, meaning that the peak lists for both negative and positive ionization modes were input. After filtering for duplicate m/z values, 1048 features were retained for analysis.

Using the BioCyc annotation library, pathway analysis identified significant enrichment in folate metabolism (folate polyglutamylation), tRNA charging, and amino acid metabolism (serine biosynthesis, arginine biosynthesis and degradation) (Figure 12). Near-significant pathways (Fisher’s p < 0.1) included amino acid metabolism (the citrulline-nitric oxide cycle and citrulline biosynthesis, aspartate biosynthesis and degradation, glutamate and glutamine degradation, asparagine biosynthesis, glycine biosynthesis and degradation, 4-aminobutyrate [GABA] degradation, proline biosynthesis, serine degradation, arginine degradation III and VI, and glycine betaine degradation), lipid metabolism (ceramide biosynthesis, triacylglycerol biosynthesis, cardiolipin biosynthesis I and II, and phosphatidylethanolamine biosynthesis), nitrogen metabolism (the urea cycle, creatine biosynthesis, and putrescine biosynthesis), nucleotide metabolism (uridine-5’-phosphate biosynthesis), folate metabolism (glutamate removal from folates), energy metabolism (glycerol-3-phosphate shuttle and glycerol degradation), fatty acid metabolism (2-methylbutyrate biosynthesis), and detoxification (glutathione biosynthesis).

**Figure 12.**
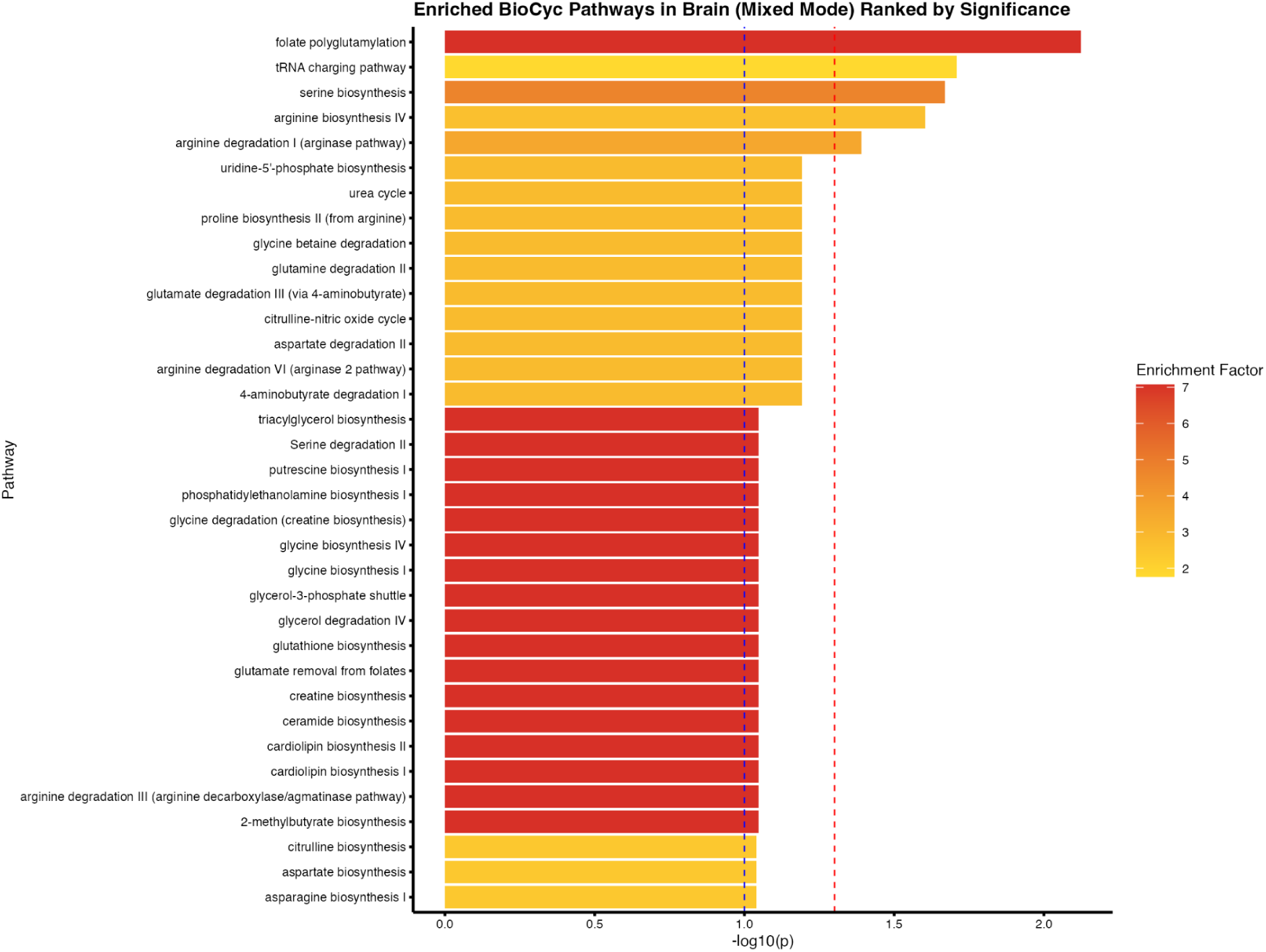
Enriched biological pathways in the mouse brain control vs. 4 hr group using the BioCyc annotation library and mixed mode. The blue dashed line denotes a p-value of 0.1 to denote near-significant results, while the red dashed line denotes a Fisher’s p-value of 0.05.

Using the KEGG annotation library, pathway analysis identified significant enrichment in amino acid metabolism (D-amino acid metabolism, arginine and proline metabolism, and arginine biosynthesis), cysteine and methionine metabolism, and xenobiotic metabolism (metabolism of xenobiotics by cytochrome P450) (Figure 13). There were no pathways that were near-significant.

**Figure 13.**
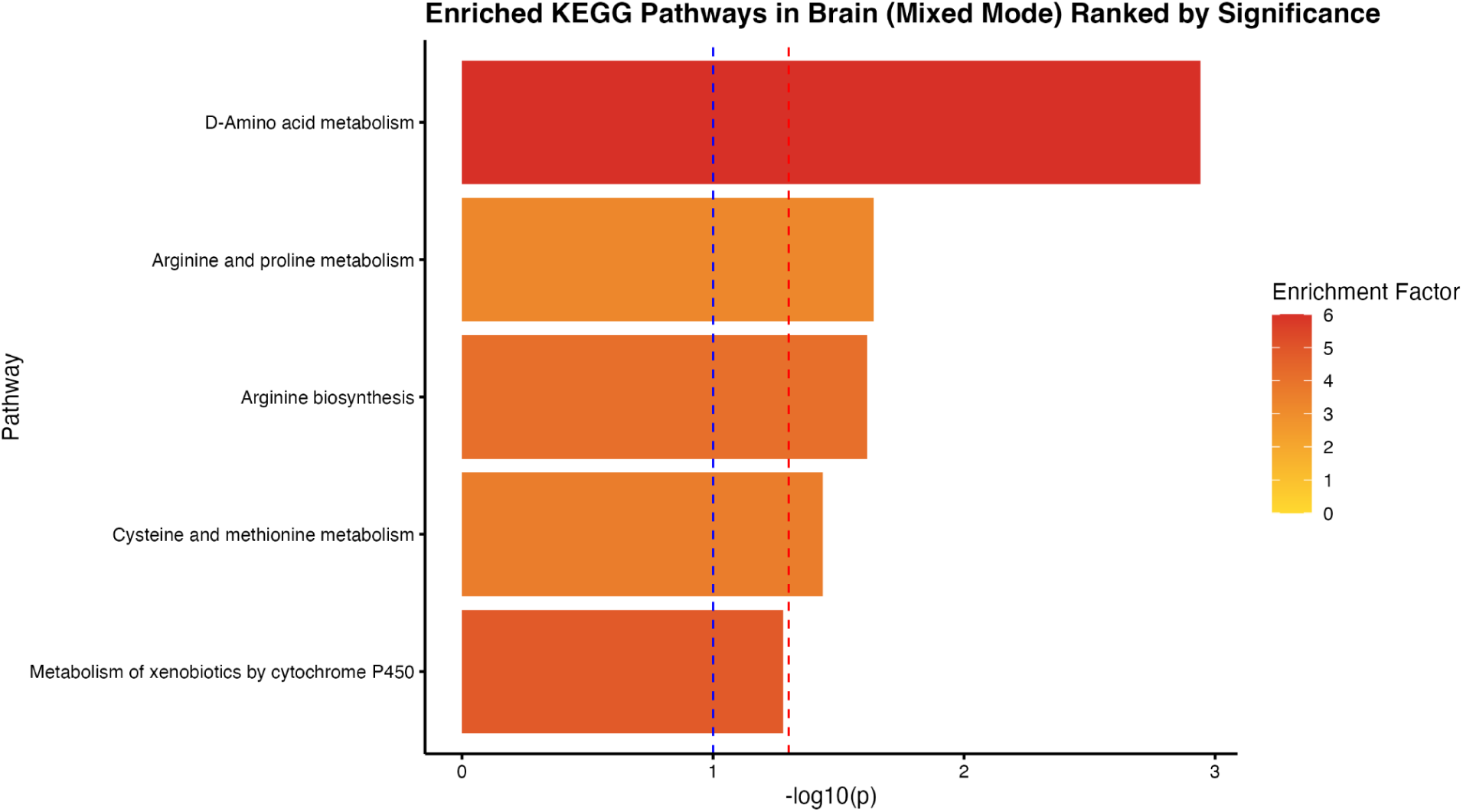
Enriched biological pathways in the mouse brain control vs. 4 hr group using the KEGG annotation library and mixed mode. The blue dashed line denotes a p-value of 0.1 to denote near-significant results, while the red dashed line denotes a Fisher’s p-value of 0.05.

Finally, we identified an additional chlorinated species that did not match our list of known TCE metabolites yet significantly differed from the control group (*p* = 1.796 × 10^-8^). As this compound had no in-house reference standard or reference library match, confidence in the identification of this compound was at level 2b (Schymanski et al. 2014). The mass spectrum of this compound exhibited prominent peaks at *m/z* 116.9072, 118.9042, 120.9012, and 122.8982. This isotopic pattern of 27:27:9:1 (M, M + 2, M + 4, and M + 6) is indicative of a trichlorinated species. Upon further probing using the SIRIUS program, which can be used for elucidating chemical structures from mass spectrum, we found that the likely chemical structure for this compound is CHCl_3_, which is the chemical formula for chloroform (Böcker et al. 2009). This was an interesting finding, as we note that chloroform is a commonly known chemical that has not been reported as a metabolite of TCE, and yet we find evidence that it may have been present in the systems of the mice who were exposed to TCE as compared to the mice who were not exposed to it (Figure 14).

**Figure 14.**
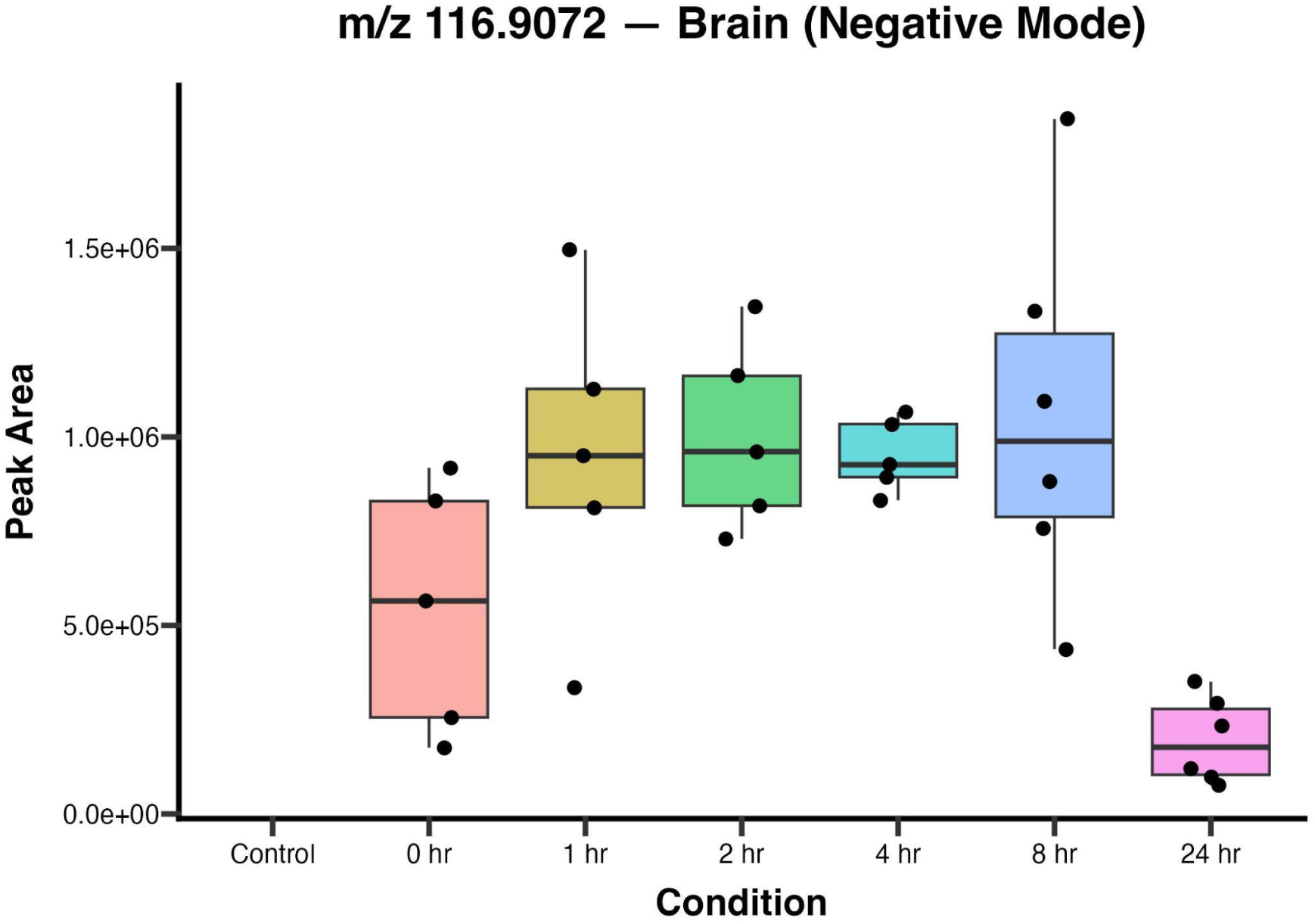
Boxplots of peak height for an unknown chlorinated compound (likely chloroform). N = 5 - 6 mice per group.

## Discussion

TCE exposure has been linked to an increased risk of Parkinson’s disease in several studies; however, TCE is highly metabolized and there is little information about which of its metabolites are present in the brain after inhalation exposure. The purpose of this study was to identify TCE metabolites in mouse brain after a human-relevant inhalation exposure. This study developed and validated a single extraction method to quantify TCE metabolites across multiple biological matrices, then identified the CYP2E1-mediated metabolites TCA, DCA, and TCEOH-gluc in mouse biospecimens following acute TCE exposure. This is the first reported study to use a human-relevant exposure paradigm in mice to assess TCE metabolite presence in the brain, highlighting how exposure route can impact chemical persistence. Here, we discuss method performance (percent recovery and matrix effects), metabolite quantities across matrices, and TCE-induced changes in global brain metabolism.

Analyte loss during extraction is a well-known limitation in the field (Zhou et al. 2003; Wong et al. 2010; Emami and Taha 2023). Percent recoveries were promising, especially in brain tissue. Plasma was the least consistent, with many concentrations showing percent recoveries well above 100%, likely due to the plasma extraction process introducing components that enhanced the signal of the compounds relative to the post-extraction spiked compounds. Our plasma extraction protocol may require modification to match performance in liver and brain.

Another common limitation in the field is matrix effects, which varied widely across the compounds, concentrations, and matrices (Cappiello et al. 2008; Trufelli et al. 2011; Panuwet et al. 2016; Cortese et al. 2020). We addressed this using a matrix-matched calibration curve.

Many studies detecting TCE or its metabolites use GC-MS for parent TCE and CYP2E1 metabolites, and LC-MS for GST metabolites (Aranda-Rodriguez et al. 2015; Luo, Hsieh, et al. 2018a). Here, we demonstrate that most of these compounds can be detected using LC-MS alone. This avoids lengthy derivatization steps required for TCA detection and enables a single extraction protocol for both CYP2E1 and GST metabolites. LC-MS also captures global metabolite information, allowing targeted and untargeted metabolomics from a single sample preparation. Our method may not provide as comprehensive metabolite coverage as a dedicated untargeted preparation, but it still enables untargeted analyses.

Within the 24-hour time course following acute TCE exposure in mice, we observed the formation and complete excretion of DCA across all three matrices. TCA and TCEOH-gluc were present at 0 hr, indicating formation during the exposure period, and were nearly depleted by 24 hr. Peak trends were similar across matrices for DCA and TCEOH-gluc, while plasma TCA peaks preceded brain and liver peaks. The early appearance of TCA in plasma likely reflects its formation during TCE exposure, whereas delayed peaks in brain and liver may reflect redistribution rather than delayed formation. TCEOH-gluc, a downstream metabolite of TCEOH, peaked early and was rapidly cleared, consistent with its role as a glucuronidated metabolite primed for excretion. No GST-mediated metabolites were detected in the brain, suggesting either no on-site metabolism of TCE by GSTs or metabolite quantities too low or transient to detect.

This is consistent with TCE literature indicating that the kidneys are the primary site of GST-mediated TCE metabolism (Lash et al. 2000; Moore et al. 2010; Luo, Hsieh, et al. 2018b). Thus, GST metabolites may have limited contribution to TCE metabolite-mediated neurotoxicity.

The AUCs indicate that exposure to TCA and TCEOH-gluc far outweighs that to DCA across all matrices. TCA and TCEOH are reported as primary metabolites of TCE, suggesting that these compounds are most likely to persist in the brain. These observations suggest that after acute TCE exposure, there is no prolonged accumulation of these compounds in the brain and that they are rapidly excreted. Further studies would be needed to assess whether these compounds accumulate in the brain after chronic exposure. If these metabolites do not accumulate in the brain after a chronic exposure, this could suggest that their effect on dopaminergic neurotoxicity is acute.

As the 4-hr time point, compared with the unexposed controls, had the greatest number of significantly different features, we used these data to identify pathways altered by TCE exposure. Many enriched pathways were consistent with prior literature. We found near-significant enrichment for increased GABA degradation, and TCE has previously been shown to affect GABA by enhancing GABA-mediated synaptic inhibition (MacIver 2009). Significant enrichment of xenobiotic metabolism by cytochrome P450s, is consistent with CYP2E1-mediated TCE metabolism. Previous work has implicated TCE as a mitochondrial toxicant, resulting in bioenergetic and oxidative stress within mammalian systems (Liu et al. 2010b; Blossom et al. 2013; De Miranda et al. 2021; Ilieva et al. 2024). While central carbon metabolism pathways were not significantly enriched, the near-significant impact on the glycerol-3-phosphate shuttle suggests early disruption to mitochondrial energy transfer (Zheng et al. 2019; Oh et al. 2024). Similarly, disruptions in amino acid metabolism are consistent with the literature characterizing TCE as an oxidative stressor (Blossom et al. 2017; De Miranda et al. 2021; Su et al. 2022; Su et al. 2023). Disruptions in cysteine and methionine metabolism are associated with oxidative stress (Travaglio et al. 2023; Martinez-Banaclocha 2023; Zhou 2023). Alterations in proline metabolism have implications for oxidative stress, redox, and energy metabolism (Plewa et al. 2021; Yao and Han 2022). Thus, acute, human-relevant exposure to TCE may begin disrupting pathways associated with chronic exposure, highlighting the rapid onset of TCE-induced metabolic dysfunction.

Other results suggest novel effects of TCE on global metabolism. BioCyc and KEGG results point to disrupted arginine biosynthesis and degradation. Arginine is an essential substrate in nitric oxide (NO) synthesis and NO has been implicated in both neuroprotection and neurodegeneration (Džoljić et al. 2015; Wu et al. 2021). Few studies have examined NO in the context of TCE exposure, though those that have reported increased NO production, consistent with disruption of arginine (Shen et al. 2007). Both libraries also indicate disruption of one-carbon metabolism via folate polyglutamylation and serine biosynthesis. Serine and folate are both key factors in one-carbon metabolism, deficiencies which have been associated with Parkinson’s disease (Tedeschi et al. 2013; Murray and Jadavji 2019). Disruptions in one-carbon metabolism can alter methylation, with downstream consequences for nucleotide synthesis and DNA methylation. TCE exposure has not been implicated in disrupted tRNA charging, yet our analysis finds this pathway enriched. tRNA charging pathways have been implicated in Parkinson’s disease, making this a worthwhile direction for future TCE research (Ko et al. 2005; Lu et al. 2014; MacMahon Copas et al. 2021; Burgess and Storkebaum 2023). D-amino acids may also play a complex role in neurodegeneration. D-serine is depleted in the cerebrospinal fluid and substantia nigra of Parkinson’s, disrupting NMDA receptor-mediated transmission (Nuzzo et al. 2019). D-tyrosine may be neuroprotective as a tyrosinase inhibitor limiting the overproduction of dopamine (Greggio et al. 2005; Park et al. 2018; Carballo-Carbajal et al. 2019). TCE’s disruption of D-amino acid metabolism warrants further investigation, as depletion of either could disrupt innate neuroprotective mechanisms.

One limitation of this study was that neither TCE nor TCEOH could be detected. TCE, as a volatile compound, and TCEOH, as a relatively small alcohol, are difficult to stably ionize and detect using LC-based methods. Detecting TCE and its metabolites could have informed formation and conversion rates, though this was not our primary focus. An alternative ionization method (e.g., atmospheric pressure chemical ionization) or derivatization of TCEOH into a readily ionizable molecule, may be necessary for its detection. Regardless, TCEOH-gluc could be considered a proxy for the detection of TCEOH, given that it is substantially converted into TCEOH-gluc (Merdink et al. 2008; Lash et al. 2014). Metabolite behavior appears to include inter-compartmental transfer, as well as formation and excretion, evidenced by mismatched metabolite trends across matrices and by the calculated transfer rate constants, suggesting that the half-lives derived from the fitted model rate constants should be interpreted cautiously.

Additional sampling time points and further optimization of plasma TCA quantification could address these issues with toxicokinetic model fit. In summary, we successfully developed a protocol to extract and detect numerous CYP2E1- and GST-mediated TCE metabolites. In particular, this protocol is useful for neurotoxicological studies, as it provides a reliable method for measuring metabolites in brain tissue. In addition, we demonstrate that an acute, human-relevant exposure to TCE results in the formation of CYP2E1-mediated TCE metabolites that can reach the brain, supporting the notion that neurotoxic effects of TCE may be metabolite-mediated. Of the detected metabolites, TCA- and TCEOH-derived metabolites appear to contribute most to overall exposure, whereas DCA is rapidly cleared and may contribute less. Global metabolomic analysis indicates that acute TCE exposure disrupts neurotransmission and oxidative stress pathways, suggesting that TCE-induced metabolic disruptions occur rapidly. Taken together, these findings support a model in which inhaled TCE is converted into metabolites that are distributed into the brain. These metabolites are likely involved in the neurotoxicity that leads to PD-related pathology. Analysis of TCE metabolites will be necessary for the field to determine the underlying mechanisms of TCE toxicity as it relates to PD. Future studies assessing chronic exposure and targeted mechanisms will help to determine the long-term consequences of TCE metabolite exposure and their relevance to human health.

## Acknowledgements

We thank Ashley Adamson, Gwendolyn Cohen, Teel Walters, and Allie Smith for their assistance with the mouse inhalation study, Brooklynn McNeil for her assistance using the TraceFinder software, Diane Re for her general feedback throughout the study, and Vrinda Kalia for her advice on the MetaboAnalyst software. Figure 1 was created using BioRender.

## Data Repository

R code and data used for this study is freely accessible on GitHub through the following link: https://github.com/jdicent1/TCE-Mouse-Metabolism

All data is accessible through Dryad using the following link: https://doi.org/10.5061/dryad.1c59zw4d0

Inhalation Chamber Exposure Protocol: https://dx.doi.org/10.17504/protocols.io.eq2lym6rwlx9/v1

