## Supplementary Information for "Metabolism of trichloroethylene following inhalation exposure in mice"

Affiliations:
[1]Department of Environmental Health Sciences, Mailman School of Public Health at Columbia University, New York, NY, United States.
[2]IARC for Carolina?
[3]Center for Neurodegeneration and Experimental Therapeutics, Department of Neurology, University of Alabama at Birmingham, Birmingham, AL, United States.
[4]Department of Molecular Pharmacology and Therapeutics, Vagelos College of Physicians and Surgeons, Columbia University, New York, New York, NY, United States.

### SI1: Standard Curve Preparation

Properties of the TCE metabolite and the final concentration of the stock solutions for each, as noted in their Safety Data Sheets (SDS), are listed in Table SI1.1. TCEOH and DCA were the only two compounds in a liquid state at room temperature; as such, their reported densities (1.55 g/mL and 1.56 g/mL at room temperature) were used to calculate the volume of each compound required to achieve the desired concentration.

| Compound | Molecular Weight (g/mol) | Weight or Volume of Compound | Volume of Solvent (µL) | Solvent Used | Concentration (mg/mL) | Concentration (mM) |
|---|---|---|---|---|---|---|
| TCA | 163.4 | 100 mg | 1000 | 0.1% Formic Acid in $H_2O$ | 100 | 611.995 |
| DCA | 128.94 | 100 mg (64.10 uL) | 1000 | Acetonitrile | 100 | 775.5545 |
| MCA | 94.5 | 100 mg | 1000 | 0.1% Formic Acid in $H_2O$ | 100 | 1058.2 |
| TCEOH | 149.4 | 100 mg (64.52 uL) | 1000 | Acetonitrile | 100 | 669.344 |
| TCEOH-gluc | 325.5 | 5 mg | 200 | 0.1% Formic Acid in $H_2O$ | 25 | 76.8049 |
| DCVG | 402.3 | 10 mg | 400 | 0.1% Formic Acid in $H_2O$ | 25 | 62.1427 |
| DCVC | 216.08 | 5 mg | 200 | 0.1% Formic Acid in $H_2O$ | 25 | 115.6979 |
| NAcDCVC | 258.122 | 5 mg | 200 | 0.1% Formic Acid in $H_2O$ | 25 | 96.8534 |

**Table SI1.1.** Preparation of metabolite stocks in acidic mobile phase. Note that the same process was repeated for each mobile phase composition tested. Only the composition of the acidic mobile phase is outlined here for simplicity.

To prepare a 2.5 mM mixed standard solution (2 mL total), the calculated volumes of each individual stock solution were combined with 178.61 µL of acetonitrile and 1546.70 µL of 0.1% formic acid in water. This resulted in a final solvent composition of 90% 0.1% formic acid in water and 10% acetonitrile. The specific volumes of each stock solution used to create this mixture of TCE metabolite standards are listed in Table SI1.2.

| Compound | Concentration of Stock (mg/mL) | Concentration of Stock (mM) | Stock Volume (µL) |
|---|---|---|---|
| TCA | 100 | 611.9951 | 8.17 |
| DCA | 100 | 775.5545 | 6.45 |
| MCA | 100 | 1058.2 | 4.73 |
| TCEOH | 100 | 669.344 | 7.47 |
| TCEOH-gluc | 25 | 76.8049 | 65.10 |
| DCVG | 25 | 62.1427 | 80.46 |
| DCVC | 25 | 115.6979 | 43.22 |
| NAcDCVC | 25 | 96.8534 | 51.62 |

**Table SI1.2.** Volumes of stock solutions for 2.5 mM mixed standard solution.

Using this stock solution, we conducted two serial dilutions in parallel, one for the 10 nM to 1 mM range and another for the 20 nM to 2 mM range. The 2 nM to 2 mM concentrations were used to spike the matrix, yielding final concentrations of 10 nM to 1 mM, while the 10 nM to 1 mM concentrations were used for neat standards. The dilutions for this are reflected in Table SI1.3. All dilutions were prepared to a final volume of 1000 µL.

| Final Concentration | Initial Stock Concentration | Volume of Stock (µL) | Volume of Solvent (µL) |
|---|---|---|---|
| 2 mM | 2.5 mM | 800 | 200 |
| 1 mM | 2.5 mM | 400 | 600 |
| 200 µM | 2 mM | 100 | 900 |
| 100 µM | 1 mM | 100 | 900 |
| 20 µM | 200 µM | 100 | 900 |
| 10 µM | 100 µM | 100 | 900 |
| 2 µM | 20 µM | 100 | 900 |
| 1 µM | 10 µM | 100 | 900 |
| 200 nM | 2 µM | 100 | 900 |
| 100 nM | 1 µM | 100 | 900 |
| 20 nM | 200 nM | 100 | 900 |
| 10 nM | 100 nM | 100 | 900 |

**Table SI1.3.** Dilutions of TCE metabolite standards for matrix-matched calibration experiments.

### SI2: Internal Standards Concentrations

Rafoxanide and spiroxamine were the inter-injection internal standards used for our experiments. 10 uL of the 1500 ppb stock of rafoxanide and spiroxamine was added to each tissue sample post-extraction, for a final concentration of 150 ppb. Since the plasma samples did not include a drying step, the 100 ppm stock of the pre-injection standards was added to acetonitrile along with the isotopically labeled internal standards stock, such that the final concentration of the pre-injection standards was 150 ppb, which would remain the same concentration of the standards throughout the entire extraction protocol.

For the TCE metabolite standard curves, 100 uL of the 1000 ppm stock was added to 1 mL of 0.1% formic acid in water and acetonitrile (90:10) to create a 100 ppm stock. This 100 ppm stock was then diluted into two separate stocks: 8.3 uL was added to 5 mL of 0.1% formic acid in water and acetonitrile (90:10) to create a 166.67 ppb stock, while 2.5 uL was added to 1 mL of 0.1% formic acid in water and acetonitrile (90:10) to create a 250 ppb stock. The 166.67 ppb stock was used to yield a 150 ppb concentration of the internal standards for all of the TCE metabolite standards except for the 1 mM stock, for which the 250 ppb stock was used. Rafoxanide and Spiroxamine were not added at the other concentrations (20 nM, 200 nM, 2 µM, 20 µM, 200 µM) because they were used to spike the blank matrix to assess matrix effects and recovery and to generate a matrix-matched calibration curve. Adding the internal standards this way resulted in a small carryover, which was calculated using a volume-weighted average of the solutions as denoted by the equation:

$$C_{final} = \frac{(V_1 \cdot C_1) + (V_2 \cdot C_2)}{V_1 + V_2}$$

The final concentrations are outlined in Table SI2.1.

| Analyte Concentration | Diluted From | IS Stock Used | Volume of IS Stock Added | Final IS Concentration (in 1 mL) |
|---|---|---|---|---|
| 1 mM | 2 mM | 250 ppb | 600 uL | 150 ppb |
| 100 uM | 1 mM | 166.67 ppb | 900 uL | 165.00 ppb |
| 10 uM | 100 uM | 166.67 ppb | 900 uL | 165.60 ppb |
| 1 uM | 10 uM | 166.67 ppb | 900 uL | 166.56 ppb |
| 100 nM | 1 uM | 166.67 ppb | 900 uL | 166.66 ppb |
| 10 nM | 100 nM | 166.67 ppb | 900 uL | 166.67 ppb |

**Table SI2.1.** IS concentrations across the TCE metabolite standard curve dilution series. Due to carryover of IS from each preceding dilution, the final IS concentration deviates slightly from the target of 150 ppb at lower analyte concentrations.

A correction factor was applied to account for this carryover in the final calculations. The correction factor was a ratio of the intended concentration ($Conc_I$) over the actual concentration ($Conc_A$), which was included in the internal standard normalization of the data in the following way:

$$\frac{Blank-Subtracted\ Peak\ Area_A}{Peak\ Area_{IS}} \times \frac{Conc_I}{Conc_A},$$

where Peak $Area_A$ is the peak area of the analyte of interest, and Peak $Area_{IS}$ is the peak area of the internal standard. This correction factor was only necessary for the TCE metabolite standard curve in clean solvent.

Isotopically labeled internal standards, used routinely by the Irving Institute for Clinical and Translational Research Biomarkers Core Lab (BCL) for metabolomics assays, were also added to the samples during preparation. Table SI2.2 provides information on the stock solution concentrations, which are diluted by adding 125 uL of stock to 5 mL of acetonitrile, and the final concentrations of these internal standards for the matrix-matched experiments.

| Compound | Stock | Working Dilution | Tissue Post Spike | Tissue Pre Spike | Plasma Post Spike | Plasma Pre Spike |
|---|---|---|---|---|---|---|
| **[13C6]-D-glucose** | 100 mM | 2.50 mM | 5.50 mM | 5.42 mM | 1.79 mM | 1.66 mM |
| **[15N]-indole** | 200 µM | 5.00 µM | 11.00 µM | 10.84 µM | 3.57 µM | 3.32 µM |
| **[2-15N]-L-lysine** | 3.00 mM | 75.00 µM | 165.00 µM | 162.64 µM | 53.57 µM | 49.82 µM |
| **[15N]-choline** | 160 µM | 4.00 µM | 8.80 µM | 8.67 µM | 2.86 µM | 2.66 µM |
| **[13C5]-L-glutamic acid** | 500 µM | 12.50 µM | 27.50 µM | 27.11 µM | 8.93 µM | 8.30 µM |
| **[13C7]-benzoic acid** | 200 µM | 5.00 µM | 11.00 µM | 10.84 µM | 3.57 µM | 3.32 µM |
| **[15N]-L-tyrosine** | 1.00 mM | 25.00 µM | 55.00 µM | 54.21 µM | 17.86 µM | 16.61 µM |
| **[15N2]-uracil** | 2.68 µM | 66.90 nM | 147.18 nM | 145.08 nM | 47.79 nM | 44.44 nM |
| **[3,3-13C2]-cystine** | 1.20 mM | 30.00 µM | 66.00 µM | 65.06 µM | 21.43 µM | 19.93 µM |
| **[trimethyl-13C3]-caffeine** | 200 µM | 5.00 µM | 11.00 µM | 10.84 µM | 3.57 µM | 3.32 µM |
| **[13C5]-L-glutamine** | 500 µM | 12.50 µM | 27.50 µM | 27.11 µM | 8.93 µM | 8.30 µM |

**Table SI2.2.** Final concentrations for isotopically labeled internal standards in the matrix-matched experiments.

Table SI2.3 provides the final concentrations of these internal standards in the real samples. Note that we were unable to perform both targeted and metabolomics runs on the plasma samples due to limited sample volume, so only final concentrations from the targeted run are reported for plasma

| Compound | Stock | Working Dilution | Tissue Samples (Metabolomics) | Tissue Samples (Targeted Run) | Plasma Samples (Targeted Run) |
|---|---|---|---|---|---|
| **[13C6]-D-glucose** | 100 mM | 2.50 mM | 1.72 mM | 5.50 mM | 1.79 mM |
| **[15N]-indole** | 200 µM | 5.00 µM | 3.45 µM | 11.00 µM | 3.57 µM |
| **[2-15N]-L-lysine** | 3.00 mM | 75.00 µM | 51.72 µM | 165.00 µM | 53.57 µM |
| **[15N]-choline** | 160 µM | 4.00 µM | 2.76 µM | 8.80 µM | 2.86 µM |
| **[13C5]-L-glutamic acid** | 500 µM | 12.50 µM | 8.62 µM | 27.50 µM | 8.93 µM |
| **[13C7]-benzoic acid** | 200 µM | 5.00 µM | 3.45 µM | 11.00 µM | 3.57 µM |
| **[15N]-L-tyrosine** | 1.00 mM | 25.00 µM | 17.24 µM | 55.00 µM | 17.86 µM |
| **[15N2]-uracil** | 2.68 µM | 66.90 nM | 46.14 nM | 147.18 nM | 47.79 nM |
| **[3,3-13C2]-cystine** | 1.20 mM | 30.00 µM | 20.69 µM | 66.00 µM | 21.43 µM |
| **[trimethyl-13C3]-caffeine** | 200 µM | 5.00 µM | 3.45 µM | 11.00 µM | 3.57 µM |
| **[13C5]-L-glutamine** | 500 µM | 12.50 µM | 8.62 µM | 27.50 µM | 8.93 µM |

**Table SI2.3.** Final concentrations for isotopically labeled internal standards in the real sample preparations.

#### SI3: Matrix-Matched Sample Preparation and Concentrations

Unexposed tissue was spiked with TCE metabolite standards before and after extraction to generate matrix-matched samples. Note that the post-extraction spiked matrix-matched samples are used as matrix-matched calibrations. Table SI3.1 provides information on the stock used to generate each concentration of the pre-extraction spiked, matrix-matched tissue samples, as well as the concentration prior to drying and reconstitution. The volume of stock used for each concentration was 15.21 µL. For post-extraction spiked matrix-matched samples, the volume of stock used was 5 µL. The stocks used to generate the final concentrations for the post–extraction spiked samples were the same as those used for the pre-extraction spiked samples.

| Final Concentration | Stock Concentration | Concentration Prior to Drying |
|---|---|---|
| 100 uM | 2 mM | 28.571 uM |
| 10 uM | 200 uM | 2.857 uM |
| 1 uM | 20 uM | 0.286 uM |
| 100 nM | 2 uM | 28.571 nM |
| 10 nM | 200 nM | 2.857 nM |

**Table SI3.1.** Pre-extraction spiked matrix-matched samples. 350 µL of the sample was completely dried, then reconstituted in 100 µL to achieve the final concentrations listed. All samples were spiked with 15.21 µL of stock.

As the plasma samples used a different extraction protocol than the tissue samples, we adjusted the volumes of the stock solution added to the unexposed plasma. The plasma samples used the same stock solutions to achieve the same final concentrations as the matrix-matched tissue samples. For the pre-extraction spiked, matrix-matched plasma samples, the volume of stock used for each concentration was 8.75 µL. For the post-extraction spiked, matrix-matched plasma samples, the volume of stock used for each concentration was 5 µL.

### SI4: Secondary Toxicokinetic Analysis with Brain as a Site of Formation

A secondary toxicokinetic analysis was done using the brain and the liver as sites of formation for the metabolites. The following equations were used for this model:

$$\frac{dC_p}{dt} = -k_{pl}C_p - k_{pb}C_p - k_{el}C_p + k_{lp}C_l + k_{bp}C_b$$

$$\frac{dC_l}{dt} = k_{liver\,form} \cdot t \cdot e^{-k_{liver\,decay}t} + k_{pl}C_p - k_{lp}C_l$$

$$\frac{dC_b}{dt} = k_{brain\,form} \cdot t \cdot e^{-k_{brain\,decay}t} + k_{pb}C_p - k_{bp}C_b$$

where $C_p$, $C_l$, and $C_b$ represent concentrations in plasma, liver, and brain, respectively; $k_{\text{liver form}}$ is a rate constant that represents formation of the metabolite in the liver; $k_{\text{brain form}}$ is a rate constant that represents formation of the metabolite in the brain; $k_{\text{liver decay}}$ is a rate constant that represents the temporal decay of formation of the metabolite in the liver; $k_{\text{brain decay}}$ is a rate constant that represents the temporal decay of formation of the metabolite in the brain; $k_{pl}$ and $k_{pb}$ are rate constants for the transfer from plasma to liver and brain; $k_{lp}$ and $k_{bp}$ are rate constants for the transfer from liver and brain back to plasma; and $k_{el}$ is the elimination rate constant from plasma.

Figure SI4.1 provides observed and model-predicted TK profiles across matrices using these updated equations. Tables SI4.1 and SI4.2 provide estimated rate constants and toxicokinetic parameters and model performance metrics using these updated equations.

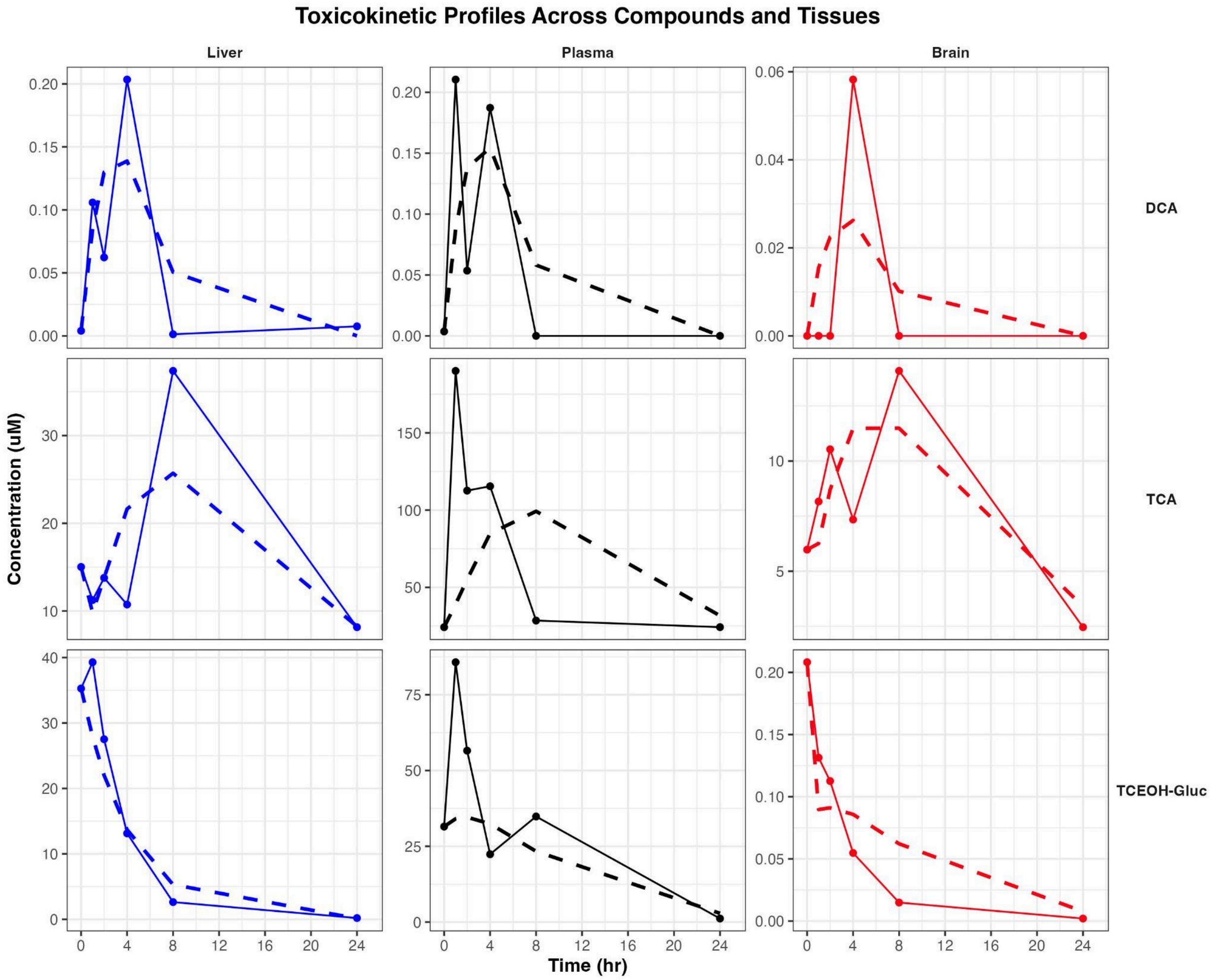


**Figure SI4.1.** Observed and model-predicted toxicokinetic (TK) profiles across matrices using the brain as a site of formation. Points and solid lines represent the observed concentration-time data, while the dashed lines represent the model-predicted concentrations.

| Compound | $k_{pl}$ | $k_{pb}$ | $k_{lp}$ | $k_{bp}$ | $k_{el}$ | $k_{liver\ form}$ | $k_{liver\ decay}$ | $k_{brain\ form}$ | $k_{brain\ decay}$ |
|---|---|---|---|---|---|---|---|---|---|
| DCA | 7.810 | 1.570 | 9.946 | 9.333 | 1.240 | 0.389 | 0.573 | 4.012 | 6.131 |
| TCA | 2.568 | 1.056 | 9.932 | 9.900 | 0.116 | 29.900 | 9.999 | 36.467 | 0.443 |
| TCEOH-Gluc | 0.000 | 0.0223 | 0.237 | 8.490 | 0.156 | 10.037 | 9.826 | 12.012 | 9.908 |

**Table SI4.1.** Estimated rate constants for DCA, TCA, and TCEOH-Gluc from the three-compartment model, including formation of metabolites in the brain.

| Compound | Tissue | Half-life (hr) | AUC | AUC ratio vs. plasma | RMSE | NRMSE | Pseudo-$R^2$ |
|---|---|---|---|---|---|---|---|
| DCA | Plasma | 0.065 | 0.854 | 1.000 | 0.067 | 0.317 | 0.440 |
| | Liver | 0.070 | 0.886 | 1.036 | 0.044 | 0.217 | 0.634 |
| | Brain | 0.074 | 0.175 | 0.205 | 0.018 | 0.303 | 0.339 |
| TCA | Plasma | 0.185 | 1196.093 | 1.000 | 73.161 | 0.385 | -0.379 |
| | Liver | 0.070 | 510.611 | 0.427 | 6.544 | 0.175 | 0.553 |
| | Brain | 0.070 | 209.601 | 0.175 | 2.302 | 0.163 | 0.596 |
| TCEOH-Gluc | Plasma | 3.892 | 611.086 | 1.000 | 23.758 | 0.277 | 0.208 |
| | Liver | 2.920 | 165.417 | 0.271 | 5.281 | 0.134 | 0.881 |
| | Brain | 0.082 | 0.734 | 0.001 | 0.030 | 0.145 | 0.823 |

**Table SI4.2.** Toxicokinetic parameters and model performance metrics for DCA, TCA, and TCEOH-Gluc across matrices in the three-compartment model that uses a formation term for brain tissue.

### SI5: Additional Information on MZmine Inputs

Below are the inputs that were used for MZmine analysis using the MZwizard tool.

Setup:

UHPLC - Orbitrap - DDA

UHPLC Module:

Check “Smoothing”
Check “Stable ionization across samples”
Crop retention time: 0.30 - 15 min
Minimum consecutive scans: 4
Approximate feature FWHM: 0.05 minutes
RT tolerance (intra-sample): 0.04 minutes
RT tolerance (sample-to-sample): 0.10 minutes

Orbitrap Module:

Select appropriate ion mode (negative or positive); Factor of lowest signal
Noise threshold MS1: 5.00
Noise threshold MS2…MSn: 2.50
Minimum feature height: 1.0E4
m/z tolerance (scan-to-scan): 0.0020 m/z or 10.0 ppm
m/z tolerance (intra-sample): 0.0015 m/z or 3.0 ppm
m/z tolerance (sample-to-sample): 0.0015 m/z or 5.0 ppm

Filters Module:

Original feature list: REMOVE
Minimum aligned samples: At least 3 samples AND 0.0%
Minimum aligned samples (in any group): At least 3 samples AND 0.0% in ANY group in condition (time point)
Uncheck “Pooled QC area RSD ≤ 20%
Check “Only keep features with 13C”
Uncheck “Apply strict shape filtering”

Annotation Module:

Check “Annotate lipids” and import desired annotation libraries

DDA Module:

Check “Apply spectral networking (FBMN/IIMN)”; other functions in this module can be done after initial analysis to improve analysis runtime.

After preparing the MZwizard, navigate to “Create batch.” Add the following step to the batch.

Feature list blank subtraction:

Aligned feature list: Those created by previous batch step
Blank/Control raw data files: Specific raw data files (select your blanks)
Minimum # of detection in blanks: 3
Quantification: Area
Ratio type: MAXIMUM
Fold change increase: 300.0 %
Keep or remove features (of rows) below fold change: REMOVE - Only keep features above fold change
Check “Create secondary list of subtracted features
Suffix: subtracted

### SI6: Additional Information on MetaboAnalyst Inputs

The data processing information for control vs. 4 hour group in brain tissue using the MetaboAnalyst functional analysis module is as follows: a total of 1195 m/z features were found in the uploaded data using a mass accuracy of 5 ppm and mixed analytical mode. The column headers of the uploaded data were m.z., p.value, t.score, r.t., and mode. A total of 136 duplicated mz features were removed, leading to 1059 input mz features retained for further analysis. The p-value cutoff for the inputted features was 1.0E-4. The Mummichog algorithm was used along with the *Mus musculus* (house mouse) BioCyc and KEGG pathway annotation libraries.

The data processing information for control vs. 4 hour group in liver tissue using the MetaboAnalyst functional analysis module is as follows: a total of 2611 m/z features were found in the uploaded data using a mass accuracy of 5 ppm and mixed analytical mode. The column headers of the uploaded data were m.z., p.value, t.score, r.t., and mode. A total of 568 duplicated mz features were removed, leading to 2043 input mz features retained for further analysis. The p-value cutoff for the inputted features was 0.001. The Mummichog algorithm was used along with the *Mus musculus* (house mouse) BioCyc and KEGG pathway annotation libraries.

### SI7: Secondary Pathway Analysis using Liver Tissue

A secondary pathway analysis was done using the data from the unexposed control vs. 4 hour group in liver tissue liver. The figures below detail this analysis, following the same structure as the brain tissue analysis presented in the main text.

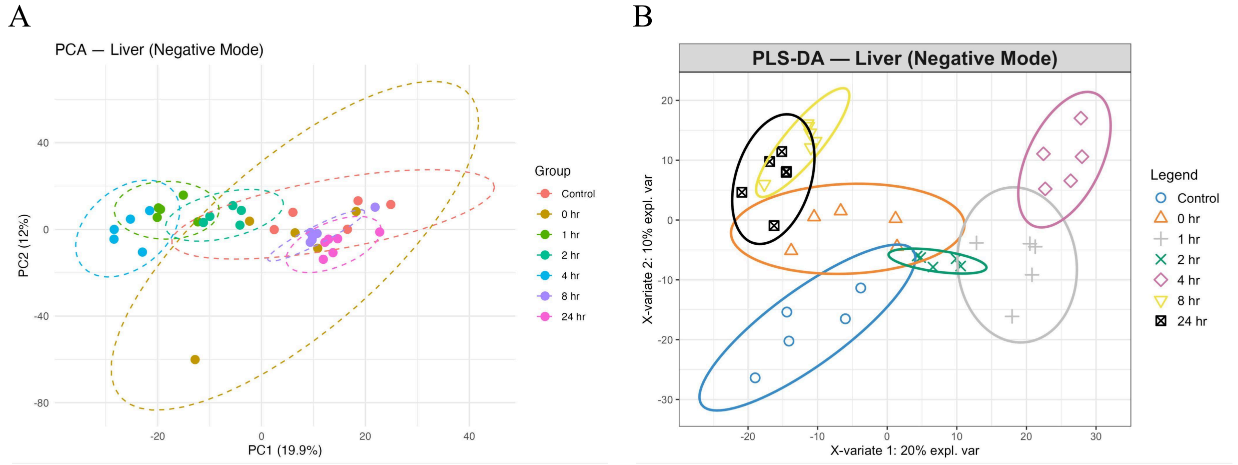


**Figure SI7.1.** PCA and PLS-DA for all groups in liver tissue in negative ionization mode. **(A)** PCA and **(B)** PLS-DA. Positive ionization mode PCA and PLS-DA graphs are omitted as there was no apparent difference between them and the negative ionization mode graphs.

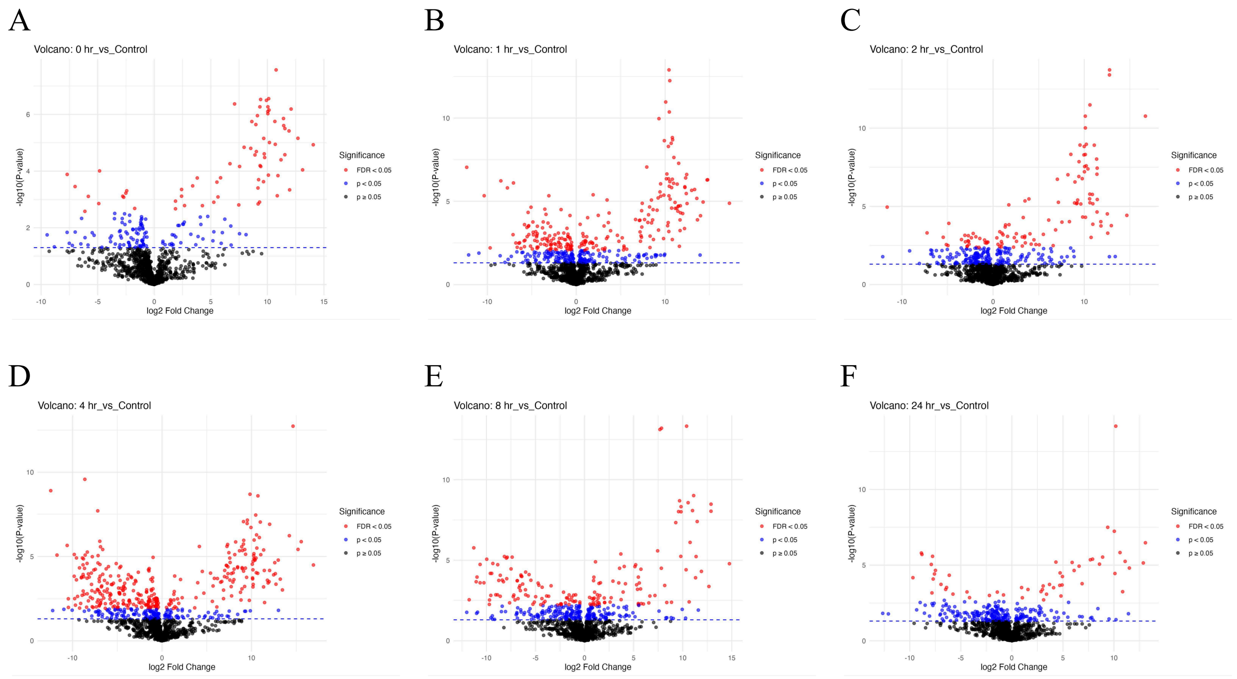


**Figure SI7.2.** Comparison of the abundances of features in the unexposed control group to the experimental groups in negative ionization mode in liver tissue. Differential abundance of control vs. **(A)** 0 hr post-exposure group, **(B)** 1 hr post-exposure group, **(C)** 2 hr post-exposure group, **(D)** 4 hr post-exposure group, **(E)** 8 hr post-exposure group, and **(F)** 24 hr post-exposure group.

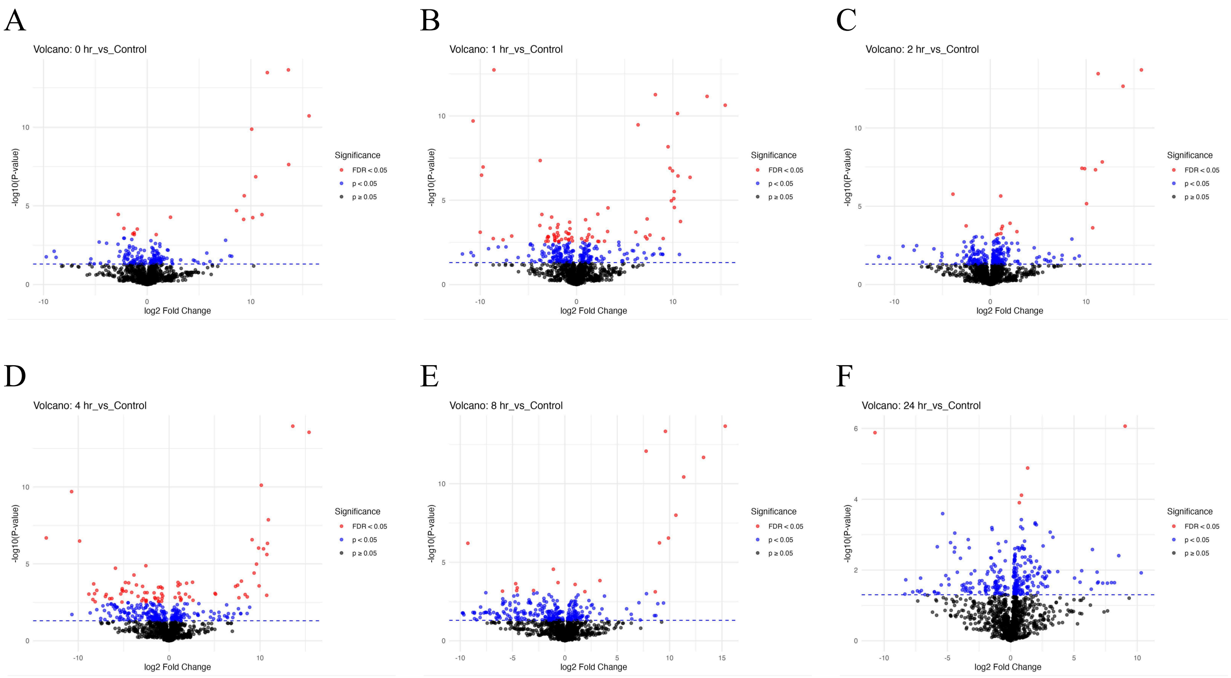


**Figure SI7.3.** Comparison of the abundances of features in the unexposed control group to the experimental groups in positive ionization mode in liver tissue. Differential abundance of control vs. **(A)** 0 hr post-exposure group, **(B)** 1 hr post-exposure group, **(C)** 2 hr post-exposure group, **(D)** 4 hr post-exposure group, **(E)** 8 hr post-exposure group, and **(F)** 24 hr post-exposure group.

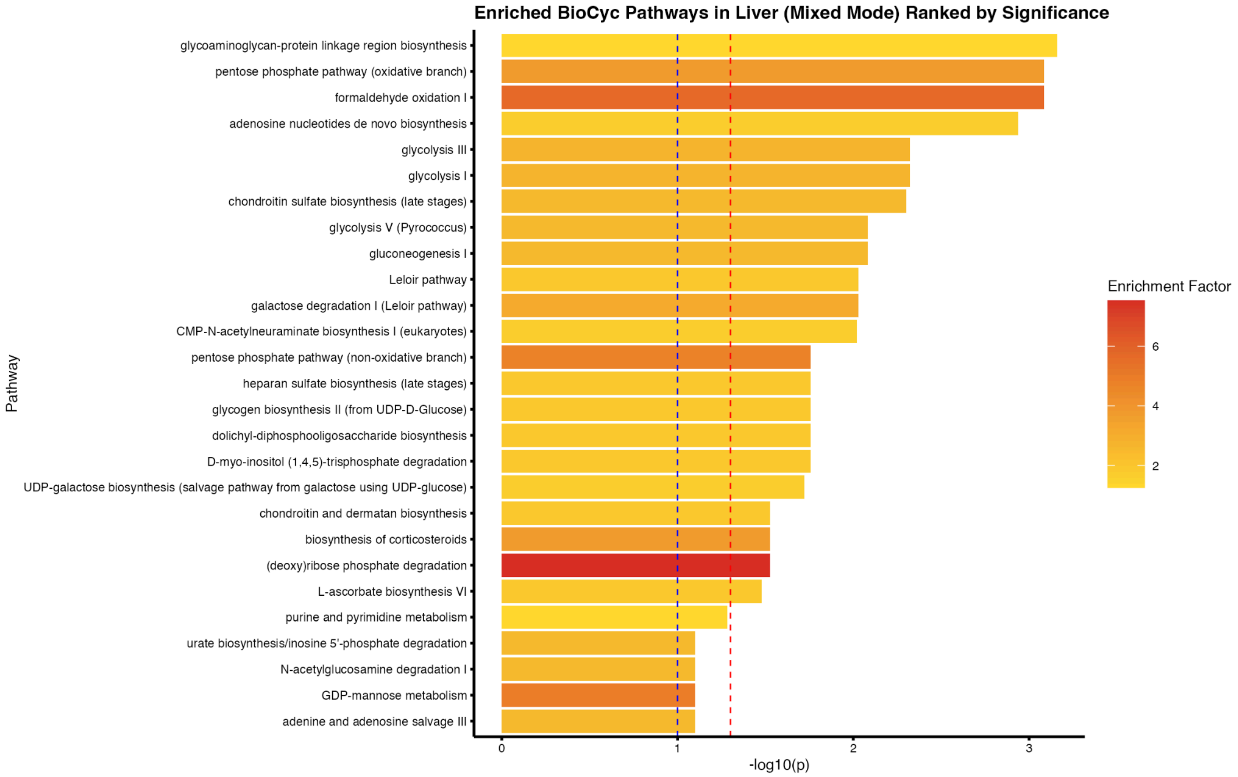


**Figure SI7.4.** Enriched biological pathways in the mouse liver control vs. 4 hr group using the BioCyc annotation library and mixed mode. The blue dashed line denotes a p-value of 0.1 to denote near-significant results, while the red dashed line denotes a Fisher's p-value of 0.05.

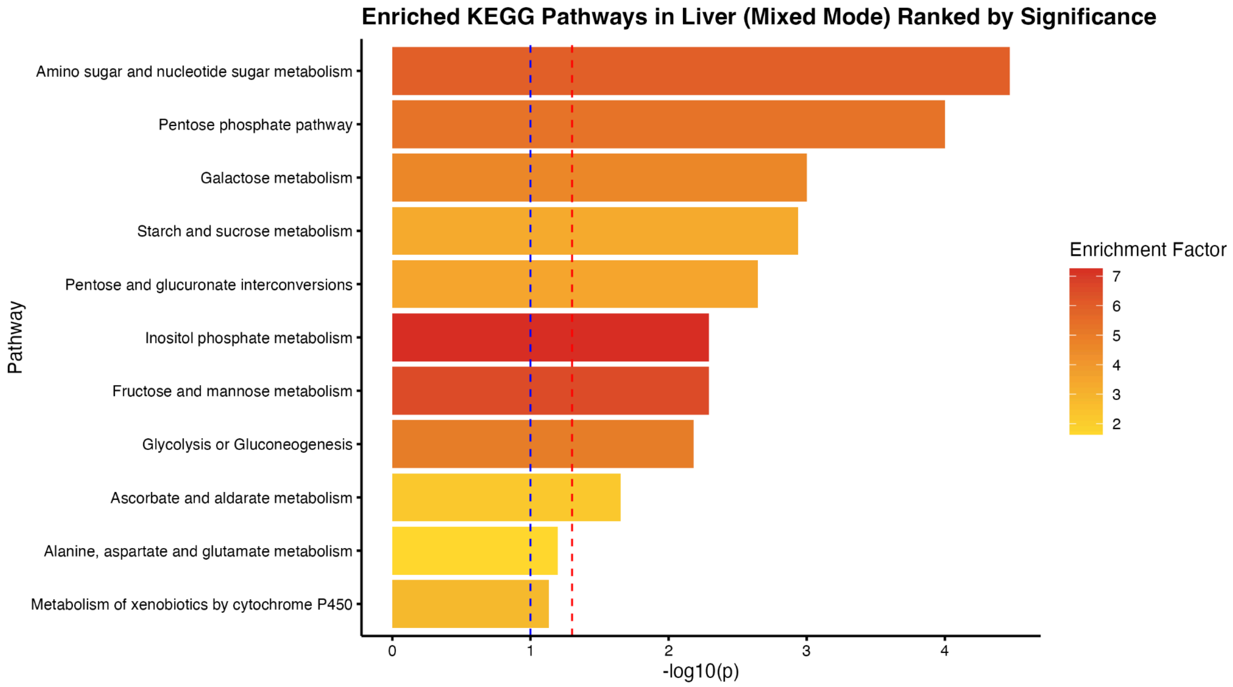


**Figure SI7.5.** Enriched biological pathways in the mouse liver control vs. 4 hr group using the KEGG annotation library and mixed mode. The blue dashed line denotes a p-value of 0.1 to denote near-significant results, while the red dashed line denotes a Fisher's p-value of 0.05.
